# In‑Situ ssDNA Isolation from dsDNA Sources as a Streamlined Pathway to DNA Origami Assembly and Testing

**DOI:** 10.64898/2026.03.19.709872

**Authors:** Enrique O. Ruiz, Kayla Neyra, Diana M. Lopez, Ruo-Wen Chen, Deepta Paramasamy, Quincy Bizjak, Patrick D. Halley, Yin Wei, Marcos Sotomayor, Michael G. Poirier, Divita Mathur, Carlos E. Castro, Wolfgang G. Pfeifer

## Abstract

Scaffolded DNA origami has become a valuable nanoscale tool for applications in biomedical and physical sciences. Critical to leveraging the modular and programmable properties of DNA origami nanodevices is access to the scaffold strand, a long single-stranded DNA (ssDNA) of precise length and sequence, which is folded into a compact shape via piecewise base-pairing with many staple strands, short ssDNA oligonucleotides. Current methods to produce and manipulate long ssDNA scaffolds can be costly, time-consuming, and cumbersome. In contrast, methods to produce and manipulate the sequence of double-stranded DNA (dsDNA) are efficient and scalable. Here, we present a method for the rapid isolation of target ssDNA sequences from a variety of dsDNA sources using oligonucleotides as blocking strands that bind continuously to the undesired strand, thereby releasing the target scaffold strand. We report successful ssDNA isolation from linear and supercoiled dsDNAs of various sequences and lengths, ranging from 769 to 15,101 nucleotides. In addition to isolating ssDNA, we demonstrated this approach enables folding of DNA origami directly from dsDNA templates using both blocking and staple strands in a single-pot thermally controlled reaction. Furthermore, we explore multi-scaffold and gene-encoding DNA origami structures, expanding the framework for application-based designs.

**Graphical Abstract:** 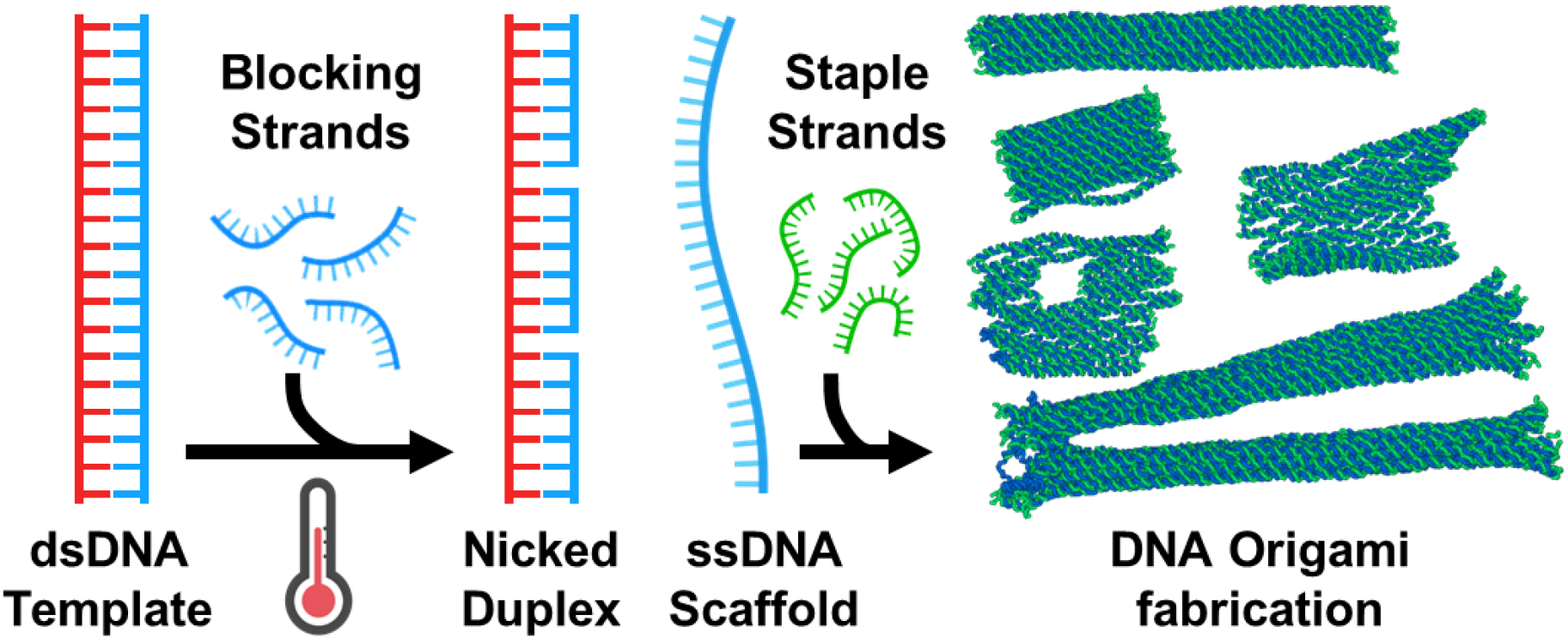

## Intro

Scaffolded DNA origami^1^ is a rapidly emerging approach that enables the design of precise and complex nanoscale shapes with sequence-specific addressability and tunable mechanical and dynamic properties^2-4^. These nanostructures have demonstrated utility for applications in gene, drug and vaccine delivery^5-8^, biophysical measurements ^9-12^, and nanoelectronics, optics and plasmonics^13-17^. A key bottleneck in the development of these nanodevices is the ability to produce long (≳ 200 nucleotides, nt) single-stranded DNA (ssDNA) with custom size and sequence, which serves as the base material for folding DNA origami structures. While a variety of methods to engineer and produce ssDNA have been developed,^18^ these processes are often complex, expensive, or have limited versatility for customizing sequence. In contrast, methods to produce double-stranded DNA (dsDNA) of custom size and sequence are well-established, highly efficient, and widely adopted through methods like molecular cloning, Gibson or Golden Gate assembly, and polymerase chain reaction (PCR)^19^. Hence, a simple and versatile approach to generate ssDNA from a variety of custom-engineered or commercially available dsDNA sources is a critical need for DNA nanotechnology. Furthermore, methods that achieve isolation of ssDNA and direct folding into DNA origami nanostructures (DOs) will improve accessibility and versatility of DO fabrication and greatly bolster the development of these nanodevices towards impactful applications.

Fabrication of DOs is based on the discontinuous hybridization of short oligonucleotides (“staple strands”, 20 - 60nts) to long ssDNA “scaffold” strands to drive folding of the scaffold into a designed compact structure^3,20^. Since the development of DNA origami folding in 2006^1^, the M13mp18 bacteriophage ssDNA genome has served as the most commonly used scaffold. Recent advances in the preparation of ssDNA have expanded the available sources, leading to scaffolds of different sizes, having orthogonal sequences, and containing target motifs for protein binding^21-28^. One powerful approach introduces the desired ssDNA sequence into a phagemid, which can then be grown through scalable biotechnological production (i.e. biomanufacturing)^21,22,24,28^. However, this approach requires significant up front effort and investment, and hence it is not practical for low-to medium-scale production, particularly when needing to test multiple scaffold sequences. Additionally, post synthesis purification measures are essential to eliminate endotoxin contamination for any downstream biomedical or cellular applications of the DO. Since dsDNA production methods are well-established, approaches to isolate ssDNA from dsDNA templates could be useful alternatives. For example, enzymatic methods, such as asymmetric polymerase chain reaction (aPCR)^23^ or lambda exonuclease digestion^29^ enable ssDNA production from dsDNA templates with low resulting endotoxin contamination, but suffer from diminishing yields for longer ssDNA (≳10,000 nt). Alternatively, dsDNA templates can be prepared with biotin or chemical modifications to anchor to streptavidin coated magnetic beads or polymer matrices, and after denaturation the anchored ssDNA strand can be separated through bead pull down, polymer precipitation, or gel separation^30-35^. However, these methods are often tedious, requiring multiple purification steps, optimization of annealing and/or denaturation protocols, or expensive or hazardous chemical modifications on the DNA, which can be challenging or even prohibitive for some labs. Thus, methods to isolate ssDNA from a variety of dsDNA sources produced either by PCR or plasmid preparation methods without the need for any DNA modifications could remove critical barriers to the broader use of custom ssDNA strands in DO self-assembly and other applications.

In addition to producing custom ssDNA scaffolds, the ability to fold DOs directly from dsDNA templates would be highly attractive to eliminate time and complexity. Folding of DOs from dsDNA templates has been demonstrated with the help of denaturing chemicals resulting in the folding of two DOs, one from each of the two strands in the dsDNA template^36^. Alternatively, other work has demonstrated folding of DOs that integrate both strands from a dsDNA template into one large structure^37^, which required deliberate design of folding pathways, larger than typical excess of staple strands, and tailored thermal annealing protocols. These efforts established an important foundation for folding DOs from dsDNA templates. However, they are still tedious relative to typical DO folding (e.g. requiring removal of chemical denaturants through dialysis) or impose design constraints (e.g. careful design of scaffold routing to control folding pathways) that limit versatility for a variety of DOs.

To overcome these challenges, here we introduce a simple method to isolate a ssDNA scaffold from a variety of dsDNA templates using short oligonucleotides (∼60 nt) that bind contiguously to the strand complementary to the scaffold, leaving the target scaffold strand accessible for isolation and purification or for direct DO folding (**Figure 1**). We refer to these short oligonucleotides as “blocking strands” since they block the undesired strand from re-annealing to the complementary scaffold strand. This process is compatible with directly folding DOs from dsDNA templates in a temperature-controlled single pot reaction with inclusion of blocking strands and DO staple strands, and the addition of thermal annealing steps.

**Figure 1:**
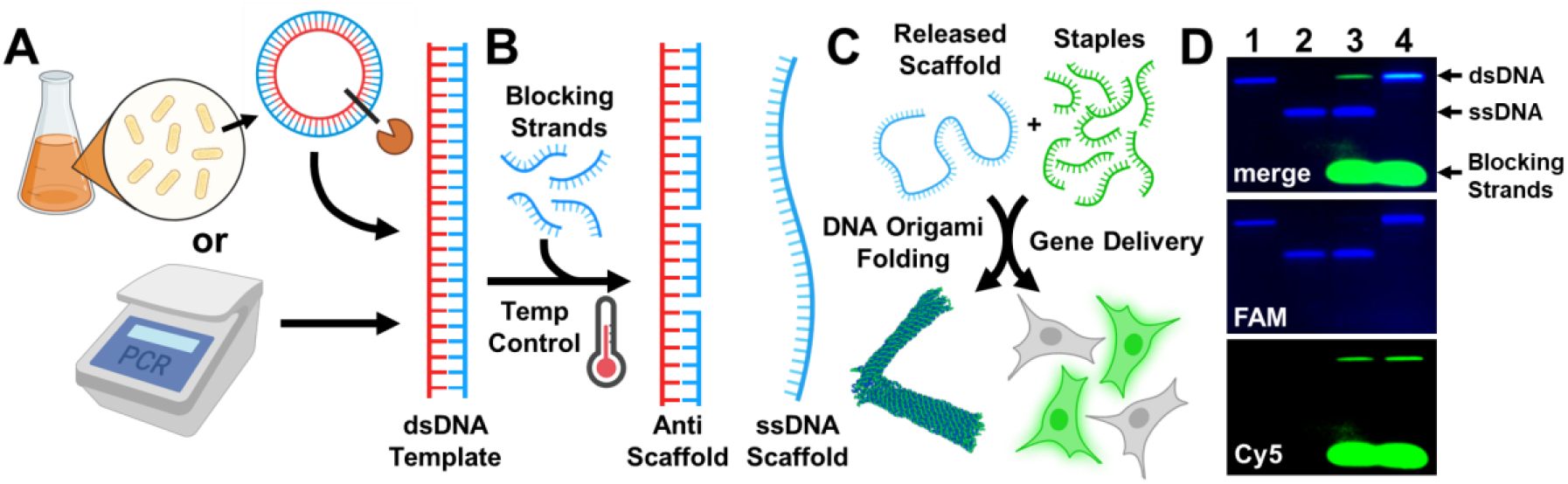
Overview of the blocking strand approach to isolating ssDNA scaffold from a dsDNA template. (A) A dsDNA template strand either in plasmid form (from bacterial prep) and enzymatically linearized or produced via PCR is (B) combined with a set of blocking strands and subjected to a thermal ramp that allows the blocking strands to bind to the anti-scaffold, thereby releasing the target ssDNA scaffold strand. (C) Schematic representing DO folding demonstrated in this work, highlighting the versatility of the blocking strands in assembling DOs of different sizes from various scaffolds, and DOs for gene delivery. (D) Agarose gel (2%) electropherograms (lane 1 = dsDNA template, lane 2 = enzymatically prepared ssDNA scaffold control, lane 3 = FAM-labeled ssDNA scaffold release through blocking strand approach, and lane 4 = blocking strand experiment sample with FAM-labeled anti-scaffold) show the migration of dsDNA template, ssDNA scaffold, and blocking strands (Cy5-labeled).

Importantly, this process does not inherently require any modifications on the DNA, making it suitable for a variety of dsDNA sources, though modifications useful for function or purification can still be used. We demonstrate the process versatility by isolating ssDNA scaffold and directly folding DOs from dsDNA templates produced by PCR as well as from biomanufactured plasmids, and we demonstrate the ability to fold a dynamic DO up to ∼15 kilobases (kb) in size. Finally, we use this approach to produce an eGFP-encoding DO and demonstrate its functionality for cellular delivery and GFP expression. Overall, our results establish a simple and versatile approach that can facilitate folding of DOs from a broad spectrum of dsDNA sources, thereby lowering barriers to the use of custom engineered scaffolds.

## MATERIALS AND METHODS

### Materials and Reagents

All DNA oligonucleotides (staple and blocking strands) were purchased from Integrated DNA Technologies (IDT) or Eurofins at 10-100 nmol scale either suspended in IDTE buffer pH 8.0 up to 100 µM or as desalted products in TE buffer pH 8. Modified staple and blocking strands (Cy5, biotin, photocleavable (PC) biotin labeled) were purchased HPLC purified and resuspended in Nuclease-Free HyPure Molecular Biology Grade Water (Cytiva, cat. #SD30538.03) to a concentration of 100 µM; concentrations were verified using a NanoDrop spectrophotometer (Thermo Scientific). The FAM labeled oligonucleotides were purchased as desalted products. The 7,249 nt M13mp18 bacteriophage genome (p7249) was produced in house as previously described^38^ and used as template for PCR to generate the 769 and 2,118 base-pairs (bp) amplicons. Lambda DNA, pUC19, and phiX174 single-stranded and supercoiled DNA were purchased from New England Biolabs (N3011S, N3041S, N3021S, N3023S). 1kb DNA ladder at 500 µg/mL (cat. #N3232S) was purchased from New England BioLabs (NEB) and the single-stranded 7K DNA ladder was acquired from PerkinElmer (cat. #CLS157950). The restriction enzymes PstI-HF (R3140S), KpnI-HF (R3142S), and PvuI-HF (R3150S) were purchased from New England Biolabs. Genetic Analysis Grade Agarose (CAS #9012-36-6), 10X Tris Borate Ethylenediaminetetraacetic acid (EDTA) (CAS #10043-35-3), Sodium Chloride (NaCl) (CAS #7647-14-5), Magnesium Chloride Hexahydrate (MgCl_2_) (CAS #7791-18-6), Tween20 (CAS #9005-64-5), and Tris Hydrochloride (Tris-HCl) (CAS #1185-53-1) were purchased from Fisher Bioreagents. EDTA (CAS #118432500) was purchased from Thermo Scientific. Dynabeads MyOne Streptavidin C1 at 10 mg/mL (cat. #65002) were purchased from Invitrogen. RealUV LED Strip Lights (cat. #PN 702.65) were acquired from Waveform Lighting. DNA Clean and Concentrator kit was purchased from Zymo Research (cat. #D4033). Information for other chemicals and supplies are mentioned in the corresponding methods sections. All sequences are provided in the attached Supplementary Tables 1-4.

### Production of dsDNA templates via PCR

PCR for 769, 2,118, 4,674, 4,845, 5,582, 7,088, and 8,013 dsDNA amplicons (ssDNA release templates) was performed (Bio-Rad C1000 thermocycler) with 250 nM primer final concentrations and 10-25 ng of the PCR template DNA (M13mp18 p7249) using the PrimeStar Max mastermix (Takara Bio). The following thermal cycler protocol was used: 1) 98°C for 60 seconds; 2) 98°C for 10 seconds; 3) 55°C for 5 seconds; 4) 72°C for 5 seconds/kb; 5) final extension of 72°C for 10 seconds. Steps 2-4 were repeated for 35 total cycles. The PCR products were then analyzed using agarose gel electrophoresis (2% agarose in 1x TBE (45 mM Tris, 45 mM boric acid, 1 mM EDTA, with 11 mM MgCl_2_, pH 8) to validate successful amplification. Once validated, the PCR products were purified using the NucleoSpin gel and PCR purification column kit (Macherey-Nagel, 740588.250) according to the manufacturer’s guidelines.

PCR for the largest dsDNA amplicons (15,101 bp ssDNA release templates for large hinge folding) was performed with 1 µM primer and 50 ng of the PCR template DNA (Lambda DNA). The PCR was carried out using LongAmp® Taq 2X Master Mix (New England Biolabs). The following thermal cycler protocol was used: 1) 94°C for 30 seconds; 2) 94°C for 30 seconds; 3) 59°C for 1 min; 4) 65°C for 15 min; 5) 65°C for 10 min. Steps 2-4 were repeated for 25 cycles. The PCR product was verified by gel electrophoresis using a 0.4% agarose gel in 0.5x TAE buffer (20 mM Tris base, 10 mM acetic acid, 0.5 mM EDTA, pH 8.0) run at 30 V for 16 hours. The amplified 15,101 bp PCR product was further verified through KpnI-HF digestion (digests Lambda DNA at two specific sites) at 37°C for one hour. The product was digested into three fragments with expected length (8,587 bp, 5,011 bp, 1,503 bp). The KpnI-HF-digested products were also verified through gel electrophoresis using a 1% agarose gel in 0.5x TAE buffer run at 80 V for 120 min.

### Production of dsDNA templates from plasmid sources

The vector pUC19 was expressed overnight using DH5α competent *E. Coli* cells in LB broth media (Fisher bioreagents) containing ampicillin (100 µg/mL). The pUC19 plasmid DNA was then extracted from the cells using QIAprep Spin Miniprep Kit (Qiagen, 27104), following the provided manufacturer’s protocol. The phiX174 plasmid was purchased from New England Biolabs. The enzymatic restriction reactions for both plasmids were carried out according to manufacturer’s guidelines to linearize template DNA. Briefly, 500 ng of DNA was added to a reaction mixture containing 5 µL of 10x rCutSmart buffer (New England Biolabs) along with 1 µL PstI restriction enzyme (20 units) and water to a final volume of 50 µL. The restriction reaction was carried out at 37°C for 30 min. The NucleoSpin Gel and PCR purification column kit (Macherey-Nagel) was used for further purification.

The GFP-encoded plasmid: pCMV-T7-eGFP (BPK1098), was acquired from the repository, Addgene (#133962). The plasmid was propagated in *E. coli* DH5α grown overnight at 37°C in Lennox broth (LB, Sigma Aldrich, cat. #L3022) and supplemented with ampicillin (100 µg/mL). Plasmid DNA was isolated using Monarch Plasmid MiniPrep Kit from New England Biolabs (cat. #T1110). The resulting DNA was eluted in nuclease-free water and quantified using NanoDrop; purity was confirmed with agarose gel electrophoresis (AGE). Plasmid DNA was then subjected to a restriction enzyme digestion with PvuI-HF using the manufacturers protocol (New England Biolabs, cat. #R3150S) to linearize the plasmid for downstream studies. After confirming complete digestion with AGE, the Zymo DNA Clean and Concentration 25 Kit (cat. #D4006) was used for further purification.

### Production of ssDNA control scaffolds

Control ssDNA scaffolds were either directly purchased or prepared using enzymatic methods. PhiX174 circular ssDNA was purchased from New England Biolabs (see Materials and Reagents). Control linear ssDNA for 769 and 2,118 nt scaffolds was prepared by exonuclease digestion using an exonuclease that selectively digests strands with a 5-prime phosphate label. The dsDNA template was produced by PCR as described above, and one primer contained a 5-prime phosphate label (Supplementary Table S2). The exonuclease digestion reaction was carried out using the Guide-it Long ssDNA Production System v2 (Takara Bio, cat. #632666) following manufacturer protocols with digestion incubation occurring at 37°C for 3 min 57 sec and 10 min 35 sec for the 769 nt and 2,118 nt scaffolds, respectively.

### DNA origami design & simulation

DNA origami nanostructures were designed in caDNAno (v2.4)^39^. Scaffold strand routings were designed to include a seam to be consistent with typical circular scaffold designs, even though linear DNAs do not require this. The extended hinge structure is based on a twist-corrected square-lattice^40^, while the other structures based on a square-lattice are not twist-corrected, and are designed to contain a helical spacing of 10.67 base-pairs. The phiX174 nanotube structure consists of a hollow cross-section with 18 dsDNA helices designed on a honeycomb lattice^41^, and eGFP-DO consists of a 36 helix-bundle cross-section also designed on a honey-comb lattice.

CaDNAno design files were converted to the oxDNA coarse-grained DNA model using tacoxDNA^41^. The resulting structures were subjected to an initial relaxation procedure to remove bond overstretching and generate starting configurations for molecular dynamics (MD) simulations. Simulations were run for 10^8^ steps with a time step of 0.001 simulation time units, using the oxDNA2 model^42-45^ in the NVT ensemble using the Andersen-like “John” thermostat with T = 20°C and a monovalent salt concentration of 1 M.

Simulations were executed using the oxDNA.org^42^ web server, which provides GPU-acceleration, as well as on computational resources provided by the Ohio Supercomputer Center (OSC). Simulation trajectories were analyzed using the python-based oxDNA analysis toolkit^46^.

All design files are available at the nanobase.org repository^47^ (2,118 monolayer: https://nanobase.org/structures/298, phiX174 nanotube: https://nanobase.org/structures/299, pUC19 rectangle: https://nanobase.org/structures/300, Lambda DNA extended hinge designs: https://nanobase.org/structures/303, M13mp18 p8064 control hinge: https://nanobase.org/structures/248, eGFP-DO: https://nanobase.org/structures/304).

### Release of ssDNA using blocking strands

Single-stranded DNA was released from linear dsDNA precursors by mixing them with a 10-fold molar excess of blocking strands (20-fold for 15kb lambda DNA), unless other excess noted. Samples were incubated at 98°C for 70 s, followed by a 5 min incubation at 64°C, before dropping the temperature to 20°C and preparing samples for further analysis or purification. Precursor concentrations of dsDNA templates were 10 or 20 nM. Release of ssDNA was verified by AGE using a 1% gel in 1X TBE buffer (45 mM Tris base, 45 mM boric acid, 1 mM EDTA, pH 8.0) run at 110 V for 90 min. Blocking strand concentrations and annealing temperatures for all cases are summarized in Supplementary Table 6.

### DNA origami folding

#### Self-assembly of DOs from control ssDNA scaffolds

Annealing of the 2,118-monolayer, the phiX174 nanotube, and the p8064-hinge was performed by mixing scaffold and staple strands in a 1:10 ratio in Tris EDTA buffer, supplemented with Magnesium chloride (5 mM Tris, 1 mM EDTA, 4-10 mM MgCl_2_). Folding was carried out with scaffold concentrations either at 10 nM (DOs based on phiX174) or 20 nM (DOs based on 2118). For folding the 2,118 monol*a*yer, the thermal annealing protocol included the following steps: 80°C for 2 min, annealing from 79-20°C at a rate of -1°C/min, and then stored at 4°C until further analysis. For folding the phiX174 nanotube, the thermal annealing protocol included the following steps: 65°C, 15 min, annealing from 64-61°C at-1°C/3min, 60°C for 5 min, 59-58°C, -1°C/10 min, 57°C for 15 min, 56°C at 25 min, 55°C at 30 min, 54°C at 45 min, 53-49°C for -1°C/60 min, 48-45°C at -1°C/42 min, 44ºC at 36 min, 43-42°C at -1°C/32 min, 41-39°C at -1°C/20 min, 38°C at 15 min, 37°C at 10 min, 36-35°C at -1°C/5 min, 34-30°C at -1°C/2 min, and then stored at 4°C until further analysis. The p8064-hinge was as folded as previously described^48^.

#### One pot self-assembly of DOs from PCR amplified dsDNA template and ssDNA release

Folding of the 2,118-monolayer from linear dsDNA was performed by first preparing a reaction mixture, containing the same components as described earlier. The thermal annealing protocols were adjusted by including an extra melting step at 98°C for 70 s, followed by an extra incubation step between 74°C and 84°C, to allow blocking strand binding and release of the target ssDNA strand.

#### One pot self-assembly of DOs from plasmid dsDNA template and ssDNA release

Folding of the phiX174 nanotube and the pUC19-asymmetric rectangle folding from corresponding supercoiled dsDNA sources requires restriction of the dsDNA into a linear fragment. We chose a restriction enzyme which shows optimal activity at 37°C and has only one restriction site in the template, such that we can include it into our thermal annealing protocol. To control its activity, we first incubate the reaction mixture at 37°C for 30 min, allowing the digestion to take place. We then continue as described earlier, with melting of all DNA species at 98°C, followed by an incubation step between 74°C and 84°C to allow release of the ssDNA. Finally, the thermal annealing protocol covers the established temperature regime for the respective DOs.

#### Self-assembly of DO from eGFP-encoding plasmid dsDNA template and ssDNA release

The pCMV-T7-eGFP plasmid (Addgene #133962) served as the dsDNA template. The ssDNA release and purification procedure is described in the SI (shown in Supplementary Figure S18). For assembly after ssDNA release, 5 nM blocking solution after biotin pull-down (Supplementary Figure S18, lane vi) and 50 nM per staple strand were combined in 1X TBE with 20 mM MgCl_2_ and Molecular Biology Grade Water. Assembly was carried out in an AnalytikJena Biometra TRIO thermocycler with the following program: 80°C for 5 min, 65°C for 15 min, 60-40°C for -1°C/h, 25°C for 25 min, and then stored at 4°C until further analysis.

### Validation and Purification of folded DNA origami by gel electrophoresis

AGE was used to validate folding of DOs, PCR products, and release of ssDNA. Agarose gels (typically 0.8-2% agarose as noted in other sections) were used (in 1X TBE: 45 mM Tris, 45 mM boric acid, 1 mM EDTA, pH 8, supplemented with 11 mM MgCl_2_ unless otherwise noted) and run for 90 min at 7.3 V/cm submerged in an iced-water bath. Loading buffer, either NEB (New England Biolabs, cat. #B7025S) or made in house (5 mM Tris, 1 mM EDTA, 10 mM MgCl_2_, and 50% glycerol), was added to samples before loading into the gel. Unless fluorophore labeled constructs were imaged, gels were pre-stained with Ethidium bromide. Gels including FAM and Cy5 labeled DNA were imaged using blue and red epi-luminescence and appropriate filters. Quantification of gel band intensities was performed using imageJ (v1.53t) or FIJI.

Purification of the 2,118 monolayer, phiX174 nanotube, and asymmetric pUC19-rectangle DOs was performed using gel extraction in which the band of interest was excised and isolated using BioRad Freeze ‘N Squeeze columns (cat. #7326165). Briefly, the agarose gel piece was added to the column and centrifuged at 13,000 g for 5 min at room temperature; the solution was collected for downstream AFM and TEM imaging. The eGFP-DO was purified via photocleavage.

### AFM imaging

DO samples were imaged with a Bruker BioScope Resolve Microscope, using the ScanAsyst in Air mode. Samples were prepared as previously described^49^ by applying 7 μL of sample to freshly cleaved mica (V1, Plano GmbH) and 2-5 min of incubation before the mica was carefully rinsed with ddH_2_O. Subsequently, we dried the mica with a gentle flow of air. Imaging was performed with triangular Silicon Nitride probes (ScanAsyst-Air, Bruker) at a typical scan rate of around 1 Hz.

### TEM imaging

Formvar/carbon-coated copper grids (Ted Pella, cat. #01754-F) were treated for 1 min using a benchtop plasma cleaner (BRANDX). Samples were prepared similar to previously described protocols^20^. Briefly, 6 µL of purified DNA origami samples were incubated on the treated grids for 8 min. Following incubation, excess sample was blotted away and the grids immediately stained using 1% uranyl acetate for 35 s. The grids were then air dried and imaged using a FEI Tecnai G2 transmission electron microscope (TEM).

### Cell culture for GFP expression study

All cell studies were carried out using human embryonic kidney (HEK293) cells (ATCC, CRL-1573). Cells were maintained in Dulbecco’s modified eagle medium (DMEM, Gibco cat. #11995-065) supplemented with 10% fetal bovine serum (FBS, Gibco cat #10082147) and 1% penicillin streptomycin glutamine (P/S, Gibco cat #15140122) in an atmosphere of 37°C and 5% CO_2_. For routine maintenance, 0.05% Trypsin-EDTA (Gibco, cat. #25300054) was used.

### Fluorescence Imaging of Cells

HEK293 cells were seeded into an Ibidi µ-well 8 channel glass slide (cat. #80826) at a concentration of 10,000 cells/well and incubated overnight to allow adherence. Transfection was performed using the manufacturer’s recommended protocol (Invitrogen, cat. #L3000001). Solution was incubated with the cells for 4 h after the transfection and then the solution was removed and replaced with complete media (DMEM, 10% FBS, 1% P/S). Cells were washed with phosphate-buffered saline 24 h post-transfection (PBS, Gibco cat. #10010-023) and stained with Hoechst 33342 nuclear dye (ThermoFisher, cat. #H3570) at a concentration of 0.5 µg/mL for 30 min. Following an additional PBS wash, live-cell imaging solution (LCIS, Invitrogen cat. #A59688DJ) was added immediately prior to imaging on a Nikon ECLIPSE Ti2 fluorescence microscope.

### Western Blot Quantification

HEK293 cells were seeded into a 12-well plate (CytoOne, cat. #CC7682-7512) at a concentration of 200,000 cells/well and incubated overnight to allow adherence. Transfection was performed using the manufacturers recommended protocol (Invitrogen, cat. #L3000001). Four hours after the transfection solution was incubated with the cells, the solution was removed and replaced with complete media (DMEM, 10% FBS, 1% P/S). Cells were trypsinized 24 h post-transfection for 2 min to obtain the cell suspension. The suspension was added to 1 mL of cold PBS, collected in 2 mL microcentrifuge tubes and centrifuged at 10,000 g for 3 minutes at 4°C to pellet the cells. Immunoprecipitation (IP) buffer comprised of 1% Triton X (Sigma Aldrich, cat. #93443), 150 mM NaCl, 50 mM Tris, 1 mM EDTA, and 1:100 protease/phosphatase inhibitor (Cell Signaling, cat. #5872S) in Nuclease-Free HyPure Molecular Biology Grade Water was prepared for cell lysis. IP buffer (50 µL) was added to the cell pellet. Samples were vortexed and incubated on ice for 45 min. Samples were frozen on dry ice for 3 min followed by an incubation on water ice for 15 min. This freeze/thaw cycle was repeated three times. Samples were centrifuged at 13,000 g for 30 min at 4°C, and the supernatant (lysate) was retained for subsequent protein quantification and immunoblot. Total protein content was determined using the Pierce BCA Protein Assay Kit (Thermo Scientific, cat. #23227) according to the manufacturer’s instructions. Absorbance was measured at 562 nm using a TECAN Spark microplate reader, and concentrations were calculated from a bovine serum albumin (BSA) standard curve. Samples were normalized to the lowest total protein concentration using IP buffer and 4X Laemmli Sample Buffer containing 10% β-mercaptoethanol (BME) (BioRad, cat. #1610747). Samples were boiled at 100°C for 5 min and equal amounts of protein (30 µg) were loaded onto a 4-20% gradient SDS-PAGE. 1X Tris/Glycine/SDS (Biorad, cat. #1610772) was used as the running buffer, and the gel was run at 117 V for 1 h 40 min. Proteins were subsequently transferred to a nitrocellulose membrane (Biorad, cat. #1620115) using wet transfer in 1X Tris/Glycine buffer containing 20% methanol at 90 V for 2 h. The membrane was blocked with intercept (TBS) blocking buffer (Li-Cor, cat. #927-60001) for 45 min at room temperature. The Rabbit anti-GFP monoclonal antibody (Cell Signaling, cat. #2956S; 1:1000) was added to the membrane and was placed on an orbital shaker at 4°C overnight. After five washes with 1X TBST (1X TBS, 0.1% Tween20), the membrane was incubated with goat anti-rabbit IRDye 800 secondary antibody (Invitrogen, cat. #SA5-10036; 1:10,000) on an orbital shaker for 1 h at room temperature. This was followed by an additional five washes with 1X TBST. To probe for β-actin, the membrane was incubated on an orbital shaker for 1 h at room temperature with Mouse anti-β-actin antibody (Invitrogen, cat. #MA1-140; 1:1000) and goat anti-mouse IRDye 680 secondary antibody (Thermo Fisher, cat. #35519; 1:10,000), followed by five washes in 1X TBST. The membrane was imaged using a BioRad ChemiDoc MP Imaging System.

## RESULTS

### Release of ssDNA from dsDNA templates using blocking strands

We propose releasing a target ssDNA scaffold strand from a dsDNA template through the use of blocking strands that are complementary to the opposite strand in the template which we refer to as the anti-scaffold. We hypothesized that separation of the two strands in the template would take place at temperatures where all or most of the blocking strands can bind, thereby releasing the target scaffold ssDNA. Keeping in mind DO folding thermal ramps begin with heating the sample to high temperatures, followed by annealing usually in the 40-60°C range, we aimed to have blocking strands that would anneal above 60°C in order to separate the temperature range of blocking strands binding from that of DO annealing. As a simple approach, we decided to test ∼60 nt long blocking strands, which showed good yield in initial tests compared to 30 nt and 90 nt analogues (Supplementary Figure S1).

To establish the approach of isolating ssDNA from a dsDNA template through the use of blocking strands, we prepared a 769 bp long linear dsDNA template (template and primer sequences in Supplementary Tables S1 and S2) via PCR (**Figure 1A**). A FAM-fluorophore label was included on either the target scaffold strand to be released or the complementary anti-scaffold strand using fluorophore modified primers. A set of 13 blocking strands that were contiguously complementary to the anti-scaffold were designed (see Supplementary Table S3 for sequences and associated melting temperatures). Additionally, we used one blocking strand that was labeled with a Cy5 fluorophore to enable separate tracking in gel electrophoresis experiments. Release experiments were carried out by mixing the dsDNA template with a 10-fold excess of the ∼60 nt blocking strands and subjecting the mixture to a thermal ramp (**Figure 1B**). The initial thermal protocol included a melting phase at 98°C for 70 s intended to separate the dsDNA template strands, followed by incubating the sample for 5 min at 64°C to allow the blocking strands to anneal to the anti-scaffold, and thus prevent it from reannealing with the target scaffold strand. The sample was then dropped directly to 20°C and then held at 4°C until further analysis. After the thermal protocol, the mixture was evaluated by agarose gel electrophoresis. We separately prepared two control samples for comparison: a positive control where a FAM labeled version of the ssDNA scaffold was prepared through an enzymatic digestion approach, and another control sample where the release experiment was run identically but the FAM label was on the anti-scaffold (details for control sample preparation in Methods). Gel electrophoresis of the release experiment sample showed a clear FAM signal with the same mobility as the positive control ssDNA scaffold (**Figure 1D**, lanes 2 and 3), while the labeled anti-scaffold control has the same mobility as the dsDNA template (**Figure 1D**, lanes 1 and 4). These results indicate that the target ssDNA scaffold strand is isolated, and the anti-scaffold binds the blocking strands to become double-stranded, effectively demonstrating the blocking strand approach. We also tested longer blocking strand incubation times, which did not lead to any significant increase in ssDNA release (Supporting Figure S2).

### Release of ssDNA is robust and achieves high yield across reaction conditions

To further study isolation of ssDNA through the use of blocking strands, we quantified the release of a target ssDNA scaffold from a 2,118 bp long linear dsDNA template (Supplementary Table S1) as a function of: 1) blocking strand annealing temperature (64, 74, or 84°C), 2) blocking strand annealing time (5 min vs 15 min), and 3) blocking strand concentration (1-fold to 25-fold excess). We again introduced a fluorophore on the ssDNA scaffold target strand through the use of a FAM labeled primer (Supplementary Table S2) to enable tracking in gel electrophoresis. Following the same overall release approach (including the melting phase and annealing phase), we incubated the dsDNA template with ∼60 nt (full list of blocking strands in Supplementary Table S3) long blocking strands that are complementary to the anti-scaffold strand within the dsDNA template and subsequently performed gel electrophoresis (**Figure 2A**). We observed a clear shift to faster mobility of the FAM labeled scaffold strand compared to the dsDNA template across all conditions, suggesting effective release of the ssDNA scaffold. At lower blocking strand concentrations (1-5 fold), we observed a small fraction of scaffold strand that appeared to remain double stranded, while most of the scaffold strand ran significantly faster suggesting it was released in ssDNA form. At these lower blocking strand concentrations, the released ssDNA scaffold ran slower than the enzymatically prepared ssDNA scaffold control (**Figure 2A**, lane 2). This is likely due to incomplete binding of blocking strands to the anti-scaffold, which allows for transient interactions between anti-scaffold and scaffold, slowing the scaffold migration into the gel. We purified and sequenced the fast-running population for the case of 1-fold blocking strands with a 5 min annealing time at all temperatures (64, 74, and 84°C), which confirmed the presence of the complete scaffold strand (Supplementary Figure S3 and Table S5).

**Figure 2:**
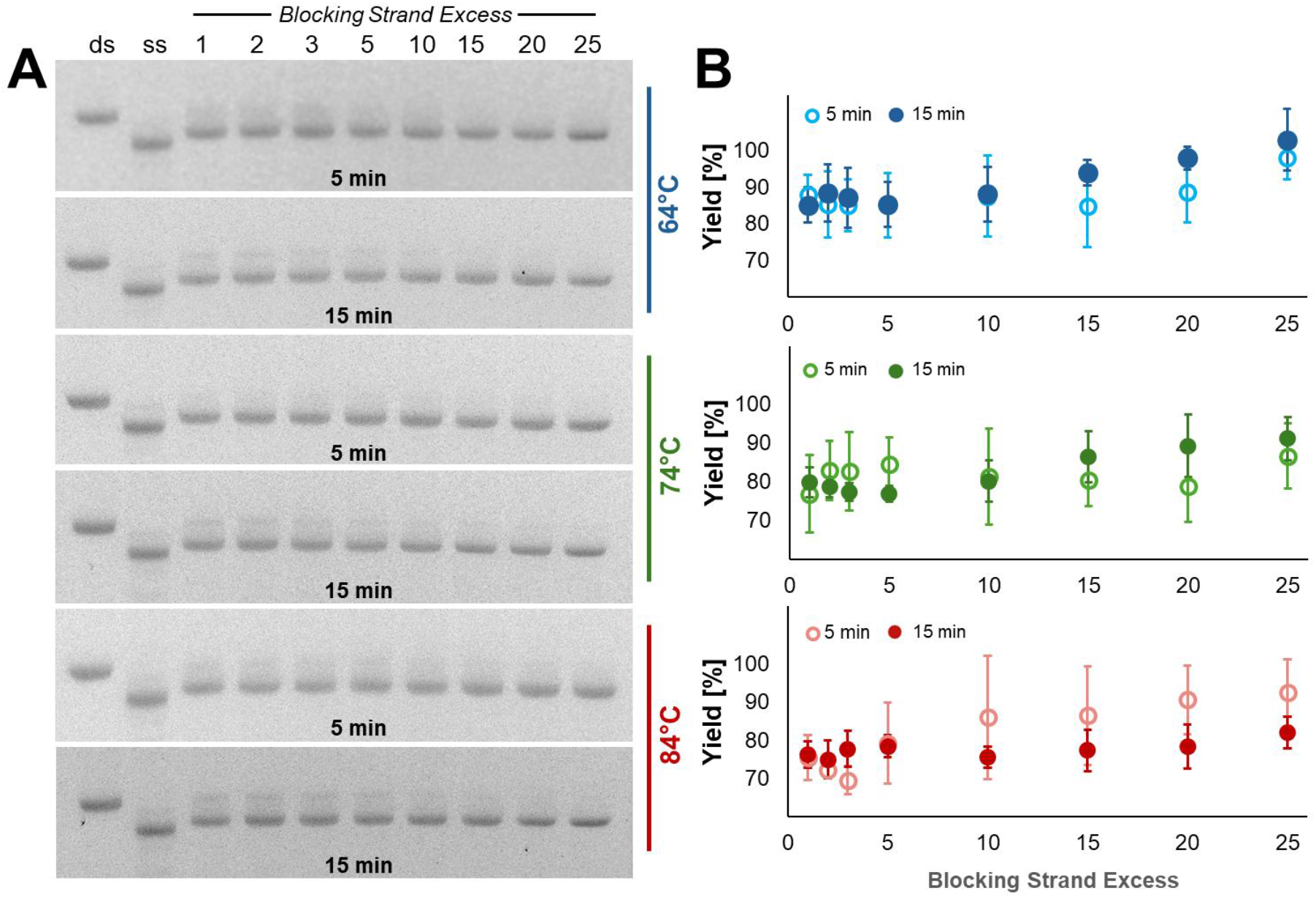
Quantification of target ssDNA release from a 2,118 bp dsDNA template under varying conditions. (A) Representative agarose gel electropherograms (1.5% in 1X TBE) of ssDNA release across 1-25 fold molar excess blocking strands (relative to dsDNA template), 5 or 15 min incubation times, and three annealing temperatures (64°C, 74°C, or 84°C). Band intensities indicate fluorescence from the FAM label on the target ssDNA scaffold strand. (B) Average ssDNA scaffold yield calculated from optical densitometry analysis corresponding to (A). Error bars represent standard deviation from the mean (n = 3).

We performed these release experiments, gel electrophoresis, and densitometry analysis in triplicate to quantify the yield of ssDNA isolation (details of yield analysis in Supplementary Figure S4). Notably, an equimolar concentration of blocking strands already led to the successful release of ∼75% (at 84°C) to ∼85% (at 64°C), with yields generally increasing with higher molar excess of blocking strands over the dsDNA template, reaching ∼85% yield or better for 25-fold molar excess of blocking strands across all annealing temperatures and times. The highest yield reached ∼100% for 64°C annealing for 15 min. This could be explained by the blocking strand melting temperatures, which are 70°C on average (Supplementary Table S3), suggesting most would stably bind at 64°C. These results show that the release of ssDNA via blocking strands is robust with even equimolar concentrations of blocking strands leading to effective ssDNA release. However, optimizing yield would require higher excess of blocking strands and selection of appropriate annealing conditions.

### Folding DNA Origami nanostructures from ssDNA released from PCR and commercial dsDNA templates

Having established the ability to release target ssDNA we then isolated ssDNA scaffolds to conduct DO folding experiments. To validate the use of isolated ssDNA for DO folding, we designed a DO structure using the 2,118 nt scaffold, optimized in Figure 2. The DO design is a rectangular monolayer with a central hole and a truncated corner as topological marker. We used coarse-grained oxDNA simulations^42-44,50^ to confirm the desired shape (**Figure 3A**). As a control sample, we fabricated the same DO with a scaffold strand generated using an established enzymatic digestion method^29^ (referred to as control ssDNA scaffold). We then used atomic force microscopy (AFM) to image both structures (prepared from either control ssDNA scaffold or released ssDNA scaffold) and observed morphologically identical structures from control (**Figure 3B** and Supplementary Figure S5A) and released (**Figure 3C** and Supplementary Figure S5B) scaffolds with both topological markers, the central hole and the truncated corner, visible and in good agreement to the oxDNA simulation. Next, we tested the blocking strand approach to produce ssDNA scaffold for a larger DO, which provides more freedom in the design space and is more comparable to commonly fabricated three-dimensional DOs. We used the phiX174 plasmid (5,386 nt), since it is commercially available as supercoiled dsDNA and as circular ssDNA. The circular ssDNA served as the control scaffold for this DO. This DO enabled us to test the release of ssDNA from a different type of dsDNA source, namely a plasmid. Prior to the release reaction, we used the restriction enzyme to linearize the supercoiled plasmid, and the ssDNA scaffold was released with ∼60 nt blocking strands and subsequently gel purified for further applications (full list of blocking strands in Supplementary Table S3, representative gel of phiX174 ssDNA release in Supplementary Figure S6). We designed and simulated the nanotube DO (**Figure 3D**), which we fabricated from both the control scaffold (**Figure 3E** and Supplementary Figure S5C) and the released scaffold (**Figure 3F** and Supplementary Figure S5D). The nanotube DO folded from the released ssDNA and exhibited conformations like those expected from simulations.

**Figure 3:**
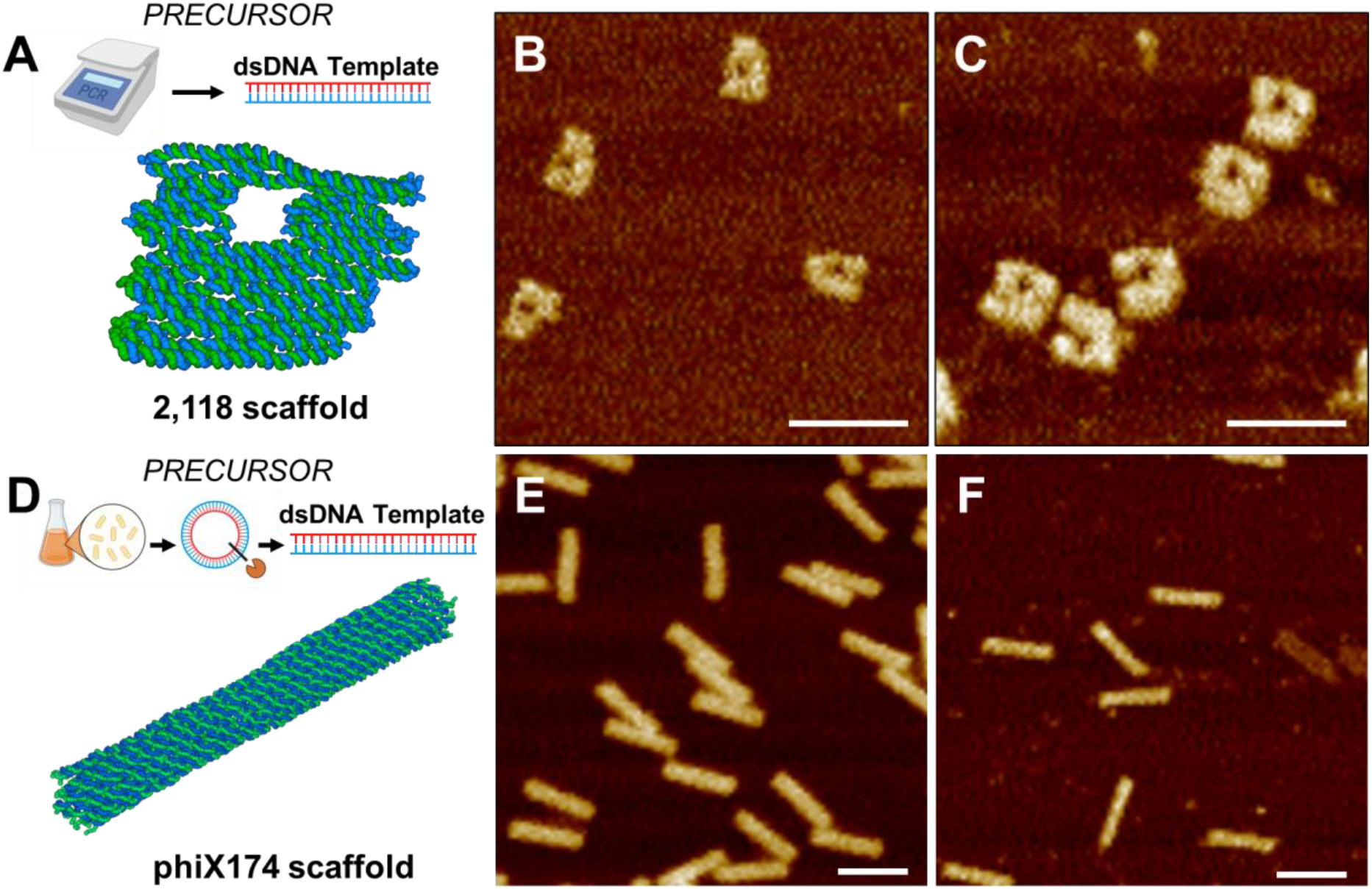
Validation of DO fabrication from scaffold isolated with blocking strand approach. (A) OxDNA simulation of a monolayer DO and the corresponding 2,118 bp dsDNA template prepared via PCR. AFM images show monolayer DO prepared using (B) the control ssDNA scaffold, and (C) the released scaffold. (D) OxDNA simulation of a 3D nanotube DO and the process by which the phiX174 dsDNA template was prepared via linearizing the precursor plasmid. AFM images show nanotube DO prepared using (E) the control ssDNA scaffold, and (F) the released scaffold. Scale bars = 100 nm.

### Folding DNA origami nanostructures in a single pot from PCR, commercial, and biomanufactured dsDNA

Since the blocking strands are designed to bind to the anti-scaffold at higher temperatures than DO annealing, we next hypothesized that the isolation of scaffold ssDNA could occur in the same single-pot reaction as folding of the scaffold into a DO (**Figure 4A**). To test this, we folded the same DOs as previously described without any intermediate scaffold purification steps. We combined the 2118 bp linear dsDNA template with its corresponding blocking strands and staple strands for the monolayer DO design. We subjected the mixture to a thermal annealing protocol, which first used 98°C to melt all dsDNA species into ssDNA components, followed by a 5 min incubation at 64°C for blocking strand annealing to the anti-scaffold, and then going through a typical DO folding protocol for staple annealing. The monolayer DO was gel-purified (Supplementary Figure S7) and then evaluated by AFM imaging. Results showed that folding directly from a dsDNA template in a single pot-reaction involving both ssDNA release and DO folding led to well-folded DO structures (**Figure 4B**, Supplementary Figure S8). Finally, we tested folding directly from dsDNA plasmid sources in a single-pot reaction using either phiX174 or the widely used plasmid pUC19. This required further increasing the single-pot complexity by integrating a restriction step to linearize the phiX174 template. We introduced a restriction digestion step at 37°C for 30 min as the first step to linearize, then increased the temperature for subsequent blocking strand and staple strand annealing steps. This ensured optimal temperature for restriction digestion and subsequent heat inactivation of the restriction enzyme. Thereafter, blocking strand binding and folding of DOs was followed by gel purification (Supplementary Figures S9-S11). AFM imaging revealed well folded DOs folded from both the phiX174 plasmid (**Figure 4C**, Supplementary Figure S12A) and the pUC19 plasmid (**Figure 4D**, Supplementary Figures S12B). TEM imaging of the three-dimensional DO nanotube also showed similar morphology of the control and one-pot structures (Supplementary Figure S13-S14). These results confirm the ability to fold directly from a variety of dsDNA sources, indicating that the more complex environment with additional components is compatible with DO folding.

**Figure 4:**
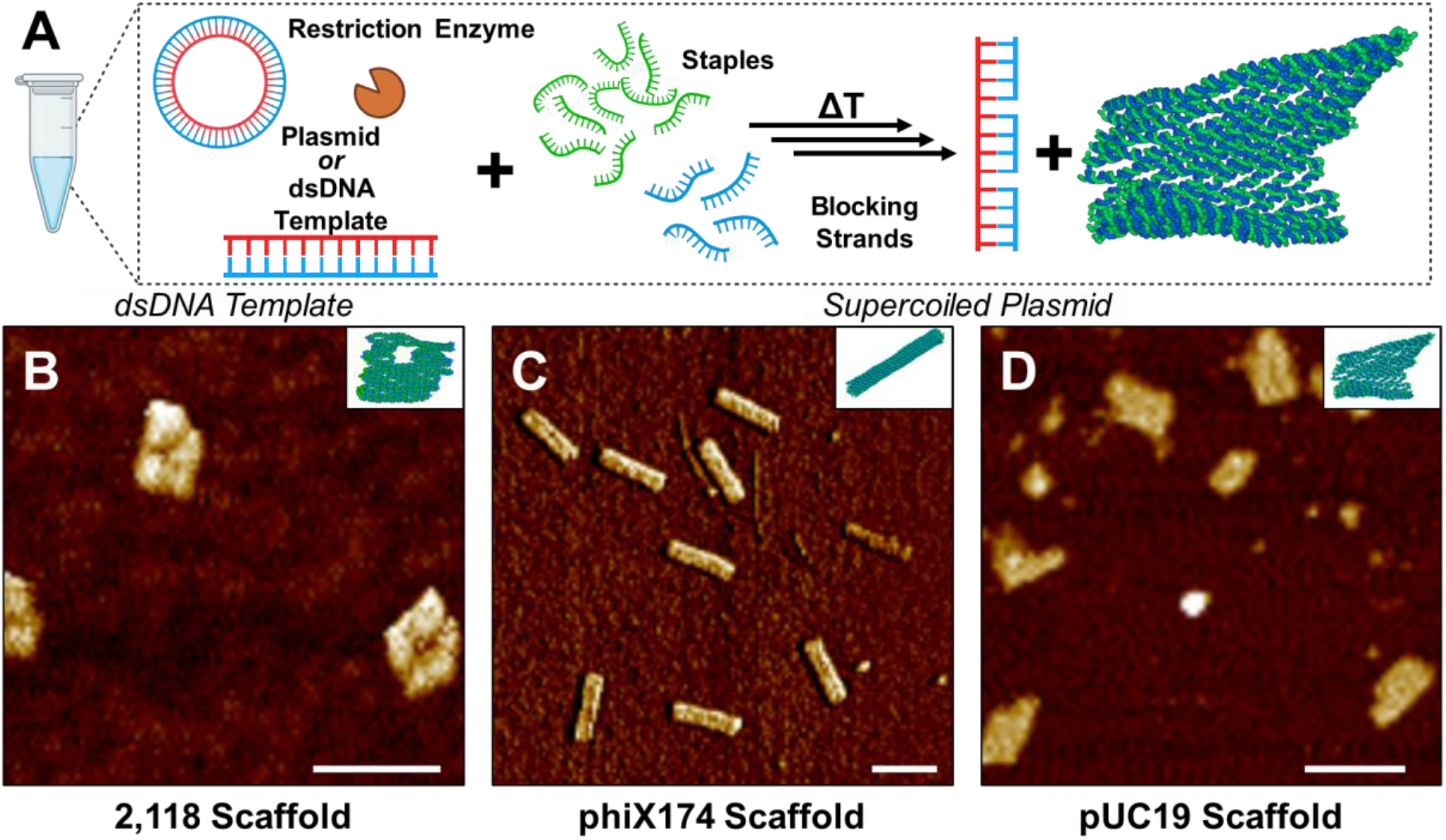
Fabrication of DOs directly from circular or linear dsDNA through a combined single-pot ssDNA release and origami assembly protocol. (A) Components for ssDNA release and DO assembly were combined in a single tube, including dsDNA, restriction enzyme (only for the plasmid sources), 5-fold molar excess of blocking strands, and 25-fold molar excess of staple strands. (B-D) Representative AFM images showing DOs assembled through the one-pot approach, (B) DO monolayer made using the 2.118 bp dsDNA template, (C) DO nanotube made using the phiX174 template, and (D) asymmetric DO rectangle made using pUC19. Scale bars = 100 nm.

### Folding a ∼15 kilo-basepair DNA origami device from one, two, or three released ssDNA scaffolds

The realization of larger sized DO devices, useful for increased structural and functional complexity, is usually achieved through hierarchical assembly routes^51,52^, which makes the fabrication more tedious. Alternatively, larger DO structures can be folded using larger scaffolds^28,53^ or multiple orthogonal scaffolds^24,54^. We sought to test whether our blocking strand approach can be used to release ssDNA up to ∼15 kb, which is approximately double the size of commonly used M13-based scaffold strands. To this end, we designed a hinge-like structure with the same cross-section as a previously published design^48,55^, but with increased arm lengths. We simulated and folded the previous hinge design using a typical M13-derived scaffold for comparison (**Figure 5A**).

**Figure 5:**
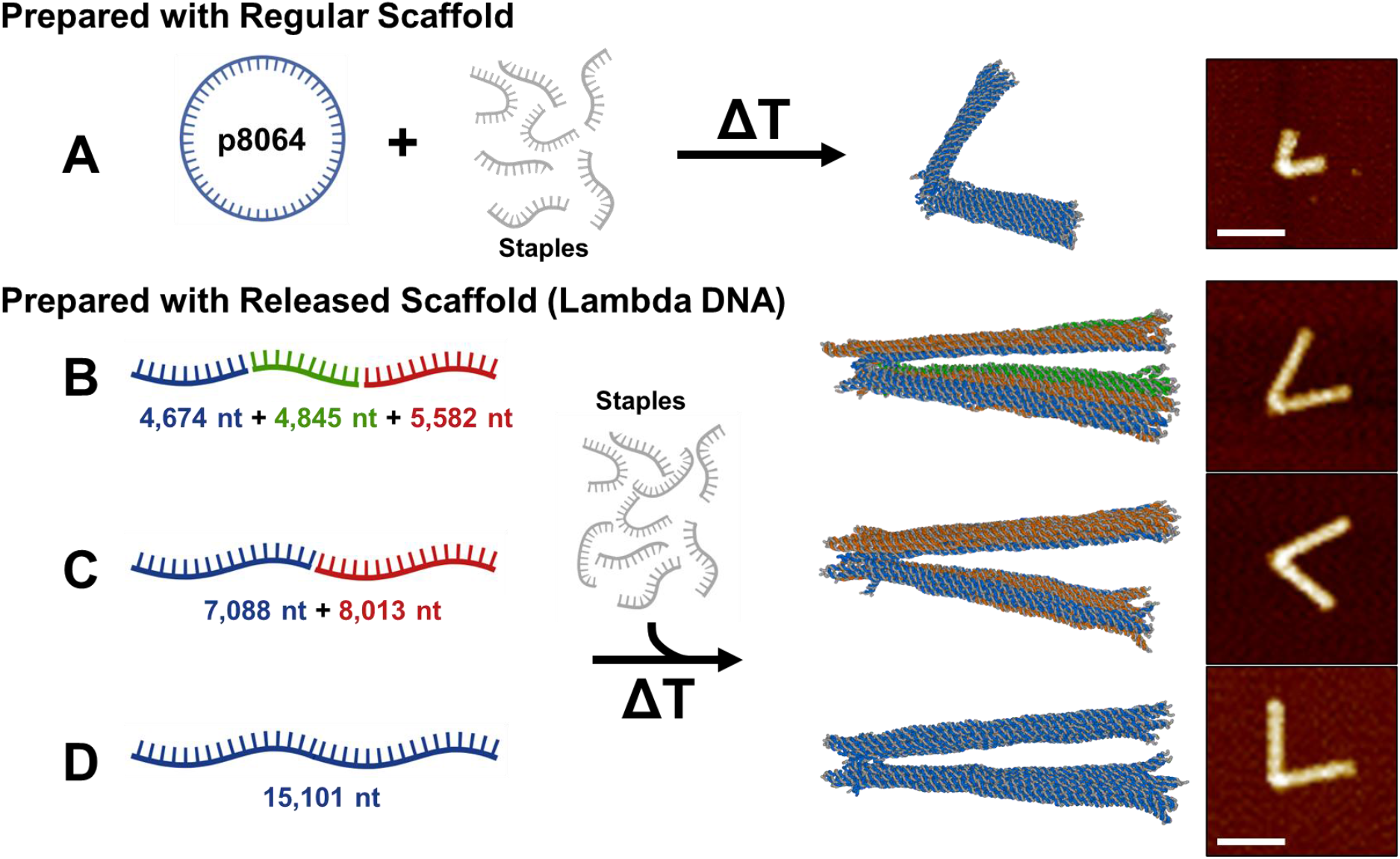
Fabrication of large DOs through release of multi-scaffolds or large scaffolds. (A) Control hinge DO was assembled using phage generated p8064 scaffold strand. Three other versions of the hinge DO were designed using scaffolds released via blocking strands from the same lambda dsDNA template, namely, (B) three orthogonal scaffolds of lengths 4,674 nt (blue), 4,845 nt (green), and 5,582 nt (red), (C) two scaffolds of 7,088 nt (blue) and 8,013 nt (red), and (D) one ∼15k nt long scaffold. For each hinge DO, the oxDNA model and a representative AFM image are shown. Scale bar = 100 nm.

To demonstrate versatility of the approach, we generated three forms of scaffolds for the extended DO hinge design from the same precursor 48.5 kb Lambda DNA. These three forms included: 1) divided into three segments (4674, 4845, and 5582 nt); 2) divided into two segments (7,088 and 8,013 nt); and 3) as one continuous 15,101 nt segment. The extended hinge DO was designed and simulated comprised of three, two, or one scaffold (**Figure 5B-D**). All desired dsDNA templates were amplified from full length Lambda DNA via PCR (template and primer sequences in Supplementary Tables S1-S2). Confirmation of the successful amplification of the target sequence was performed by restriction digestion with KpnI (restriction sites highlighted in Supplementary Table S1) and subsequent gel electrophoretic mobility analysis, yielding three fragments of the expected size (Supplementary Figure S15). We then incubated the set of three, two, or one dsDNA amplicons with corresponding blocking strands (Supplementary Table S3) and evaluated ssDNA release by gel electrophoresis (Supplementary Figure S15-16). To visualize the release of the 15,101 nt scaffold, we labeled the scaffold strand with a fluorophore using a FAM-labeled primer in the PCR, and we used one blocking strand with a Cy5 fluorophore. This provided orthogonal fluorescent signals to identify individual species in gel electrophoresis (Supplementary Figure S16).

After releasing and gel-purifying the individual ssDNA fragments, we used them in subsequent DO folding reactions. Folded DO structures were purified to remove excess staple strands using PEG precipitation^56^, and we imaged purified DOs by AFM. AFM images showed structures in the expected dimensions and in good agreement with coarse grained simulations (**Figure 5B-D** and Supplementary Figure S17). These results confirm that the blocking strand approach provides a useful route to larger DO structures either by releasing larger scaffolds or by releasing multiple orthogonal scaffolds and folding a multi-scaffold DO design.

### Fabrication and verification of GFP encoding DNA origami from a plasmid dsDNA template

Finally, we demonstrated isolation of ssDNA scaffold from a functional plasmid precursor. In recent years, DOs assembled using gene-encoding scaffold strands are being developed for gene delivery applications^5,57,58^, which retain significant advantages over traditional gene delivery vectors such as adeno-associated viruses or lipid nanoparticles^59^. Notably, DO offers compaction of the gene template, which can be beneficial for cell and nuclear entry, and a uniquely addressable set of staple strands that can be selectively modified for enhanced programmability^60-62^. Typically, gene-encoding scaffolds are obtained either through phagemid production, which is time-consuming and often yields endotoxin-contaminated ssDNA^22^, Typically, gene-encoded scaffolds are obtained either through phagemid production, which is time-consuming and often yields endotoxin-contaminated ssDNA^22^, or via aPCR, which relies on costly consumables, such as polymerase enzyme, to achieve sufficient yield.^18^ We hypothesized that our blocking strand approach could enable facile production of gene-encoded scaffold strands for DO-based gene delivery in a reagent-efficient manner.

To that end, we designed a 50 nm rectangular cuboid DO folded from a scaffold that encodes enhanced green fluorescent protein (eGFP). Importantly, crossovers along the promoter of the gene were minimized, and the scaffold was routed to position this region on the exterior of the DO, which has been shown to improve downstream transcription by RNA polymerase II. The cuboid DO was simulated with oxDNA, revealing the promoter domain on the surface that appears more accessible (**Figure 6A**). The scaffold strand was generated from a 4,119 bp plasmid precursor, which was linearized using PvuI-HF, then incubated with 5-fold molar excess blocking strands (see Supplementary Table S3 for sequences and melting temperatures). Notably, to demonstrate the compatibility of blocking strands with various downstream purification techniques, three blocking strands (two complementary to the ends and one to the middle of the anti-scaffold) were biotin functionalized to allow removal of the anti-scaffold using streptavidin-coated magnetic bead pulldown instead of gel extraction (as done in Figures 3 and 5). Affinity based purification leveraging streptavidin coated magnetic beads is often adopted due to its scalability and because it circumvents the need to expose the scaffold strand to 302 nm UV light (UV-B)^63^, which is typical for gel extraction-based purification. Here the anti-scaffold was successfully removed via magnetic beads, leaving behind a solution containing the desired eGFP-encoding scaffold and residual blocking strands (Supplementary Figure S18).

**Figure 6:**
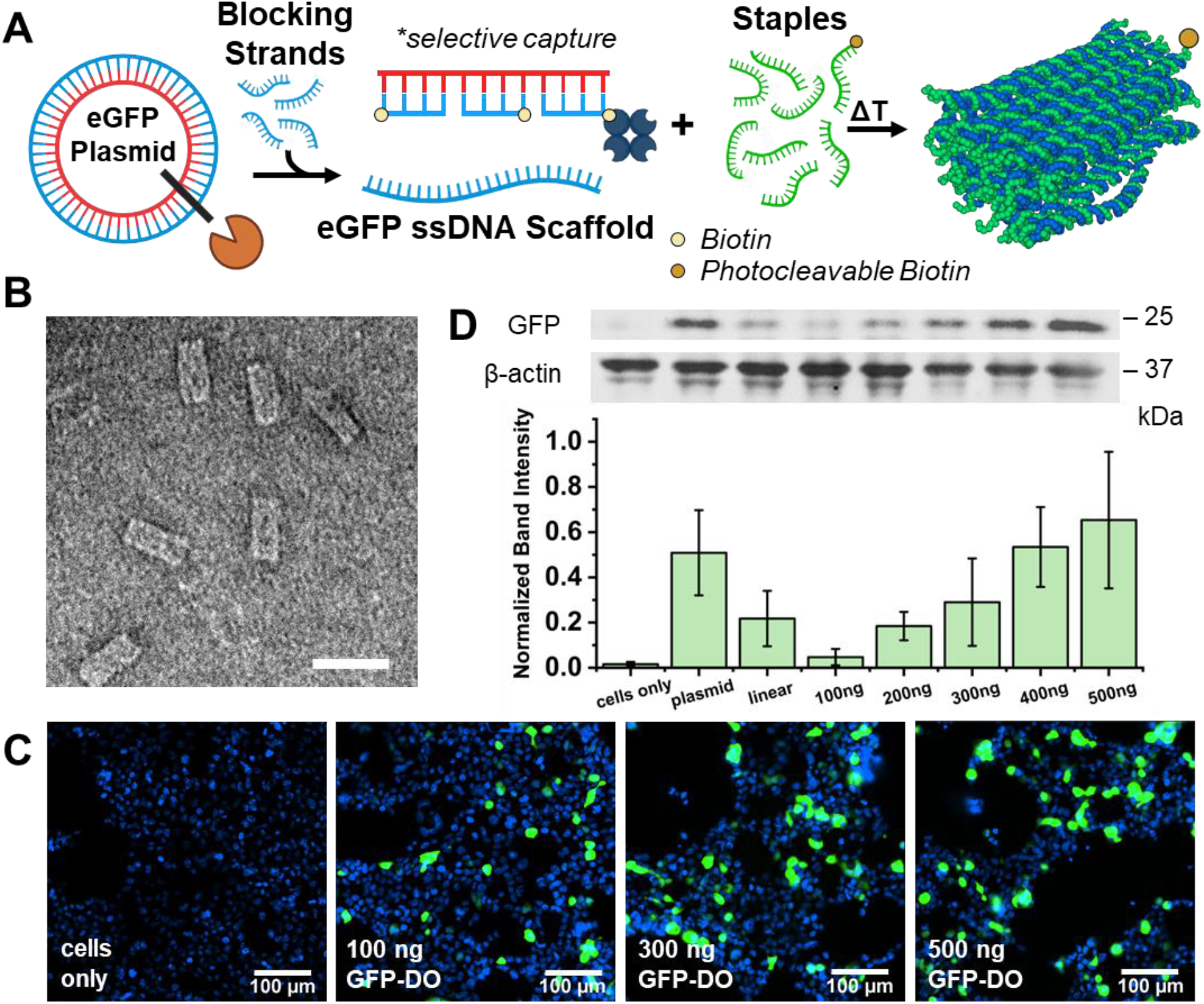
Folding and characterization of the eGFP-DO. (A) Schematic representation of the modified ssDNA release technique and subsequent DO assembly and purification. (B) TEM image of the eGFP-DO. Scale bar = 50 nm. (C) Fluorescence microscopy images of HEK293 cells 24 h after transfection with Lipofectamine3000; Blue = Hoechst nuclear stain (ex: 385 nm, em: 461 nm), Green = eGFP (ex: 475 nm, em: 520 nm). Scale bar = 100 µm. (D) Quantification of eGFP protein using western blot; samples were normalized to the housekeeping protein β-actin. [eGFP-DO] = 100-500 ng. Error is standard deviation from mean (n = 3).

This solution was then combined with 10-fold excess staple strands and thermally annealed to assemble the eGFP-DO (see Materials and Methods for details). Once again, to adopt an affinity-based purification method, we functionalized one of the staple strands with a photocleavable biotin according to a previous report^64^. Well-formed DOs were pulled out (and possibly some of the residual biotinylated staple strand) of the annealing mixture, leaving behind undesired agglomerates formed by the residual blocking and staple strands as well as excess free oligos. The eGFP-DO bound to the magnetic beads was subsequently released into a fresh vial by photocleavage of the biotin linker.^64^ Agarose gel electrophoresis (Supplementary Figure S18) and TEM (**Figure 6B**) confirmed the correct formation of the DO.

To evaluate if the scaffold generated from the ssDNA release retained its biological function, we assessed the expression capabilities of the eGFP-DO in HEK293 cells. Using lipofectamine-based transfection, we observed a dose-dependent relationship between the DO treatment and the eGFP protein expression as visualized by fluorescence microscopy and quantified via western blotting (**Figure 6C**, Supplementary Figure S19). Western blotting results showed the eGFP-DO could yield similar eGFP expression to the linear and plasmid forms of the gene, albeit at higher concentrations (**Figure 6D**, Supplementary Figure S20). These results support the hypothesis that this ssDNA isolation method can produce customized scaffold sequences that preserve the functionality of the gene of interest, demonstrating the suitability of this approach for downstream gene delivery applications.

## DISCUSSION

This work establishes blocking strands as a method for ssDNA isolation and direct DO folding from a variety of dsDNA templates. The yield of releasing target ssDNA from dsDNA was significant across a range of conditions down to 1-fold molar equivalent of blocking strands. We found improved yields when 5-fold molar excess or more blocking strands are used at an appropriate annealing temperature, reaching as high as ∼100% ssDNA release yields.

Importantly, this approach is well-suited for simultaneously isolating ssDNA scaffold strands and directly folding these DOs through strategic thermal protocols in a single pot reaction. Lastly, we broadened the scope of blocking strands to generate large ssDNA scaffolds to enable fabrication of large DOs (up to ∼15 kb) and a biologically functional scaffold encoded with GFP for gene delivery and expression. In particular, our results can facilitate the modular use of custom scaffolds in larger DO designs. Prior approaches have made tedious efforts to generate larger scaffolds^53^, multiple orthogonal scaffolds^24^, or folded from two strands of the dsDNA template^37^. We demonstrate a simple alternative approach to realize larger than usual DOs even when only smaller dsDNA sources are available. Herein, the same large hinge DO architecture was realized by a single large ∼15 kb scaffold or up to 3 unique ∼5 kb scaffold strands. Although in this case, all scaffolds in the multi-scaffold design were derived from the same original source, but it is likely that a variety of dsDNA sources could be used, increasing versatility and modularity of large DO fabrication.

There are several advantages of this blocking strand approach compared to other existing methods^18^ to generate target ssDNA strands. No chemical modifications on individual blocking strands are required. However, the technique is compatible with such modifications (e.g., biotinylation) if needed, and although not required, fluorophore modification is useful in optimizing yields. Preparation of dsDNA templates relies on widely used standard approaches such as PCR or restriction/nicking of DNA. Once a linear dsDNA template is available, the ssDNA isolation is enzyme-free, eliminating the need for meticulous experimental conditions that maintain enzyme viability and typical optimization of enzyme reactions. Most importantly, the dsDNA template could be from a variety of sources (plasmids, *de novo* gene fragments, or PCR amplicons). With plasmid being one of the most affordable and widely available dsDNA sources, the ability to produce DOs in a single pot reaction from a plasmid template is particularly relevant in streamlining fabrication, cutting down production hiccups, and facilitating scale up. These factors make the blocking strand process highly accessible and versatile. We also showed that the technique is compatible with different downstream DO purification methods, namely gel electrophoresis, centrifugal PEG purification, and affinity-based pulldown. For the latter, specific blocking strands were easily modified with biotin tags, thus exemplifing the modular nature of this approach, which is advantageous since DO applications often require choosing from a library of purification methods^7^ and added chemical modifications^16^.

While this work successfully applies the blocking strands in generating ssDNA for DO folding, we note a few limitations and directions in which further technological advancements can be made. Each dsDNA template requires a unique set of synthetic blocking strands, which effectively doubles the cost for fabrication of a particular DO design and is increasingly expensive for longer templates. However, the same set of blocking strands could be used to fold a variety of DO designs from the same scaffold. The current blocking strand set completely binds to the anti-scaffold, but future work could focus on identifying minimal sets of domain-specific blocking strands that do not span the entire anti-scaffold sequence and thus bring the cost down, ideally without compromising yield. Potentially, more advanced bioinformatics tools could help in designing a minimum set of blocking strands against homologous plasmid templates, thereby further reducing the purchasing burden. The annealing protocol for ssDNA release also requires optimization for each dsDNA template and blocking strand set since it depends on the GC content and length of the template. However, our results showed good yields (≳75%) can be achieved without any optimization effort. We demonstrate purifying the DOs via gel extraction and affinity-based pulldown, in some cases with multiple purification steps, noting that certain downstream applications could be inhibited by the nicked dsDNA byproduct (the anti-scaffold blocking strand duplex) or varying agglomerates of blocking strands and staple strands in single-pot folding. We also note that the blocking strand approach is not as scalable as phagemid-based production, which may be more suitable to produce massive amounts of a single finalized scaffold product^21,22,24,28^.

Beyond impacting the field of DNA nanotechnology, the blocking strand approach potentially advances ssDNA production for other applications as well. One notable example is the area of gene editing wherein ssDNA templates are shown to have advantages relative to dsDNA, such as reduced cytotoxicity^65,66^. Given that enzymatic and *de novo* methods have diminishing efficiency of ssDNA synthesis longer than 3,000 nt, the blocking strands could push the advancements in CRISPR-based therapeutics as well^67^. In addition to the near-term potential improvements discussed above, exploring synergies with other established techniques can further broaden the applicability of blocking strands. For instance, large-scale production of oligos using bioreactor-scale phagemid synthesis or chip-derived oligo pools has emerged as an alternative to low-cost production of staple strands^21,53^. The same frameworks could be adapted for low-cost production of blocking strands or combined sets of blocking and staple strands. Furthermore, this approach could be synergistic with other efforts to incorporate other functional elements, such as aptamers, into DO scaffolds^25^, or provide a framework for the modular use of these high-value scaffolds in multi-scaffold DOs.

## Data Availability

The experimental data sets are either included in this submission, the supplemental information, or are available from the authors upon request.

## Acknowledgments

AFM imaging was supported by the National Institutes of Health [1S10OD025096-01A1 to Gunjan Agarwal]. We acknowledge resources from the Campus Microscopy and Imaging Facility (CMIF), The Ohio State University and from the Cleveland Center for Membrane and Structural Biology (CCMSB), Case Western Reserve University. We thank members of the Castro, Poirier, Mathur, and Sotomayor labs for valuable feedback.

## Author Contribution

WGP, CEC, EOR, PDH conceived the study. WGP, EOR, KN, DML, DP, QB, YW conducted the experiments. WGP, CEC, DM, and MGP supervised experiments and gave regular feedback on experiments. MS supervised the bacterial growth for the pUC19. WGP, KN, DM, EOR & CEC wrote the paper, with feedback from all others. All authors participated in the review and editing of this manuscript. All authors have read and agreed to the published version of the manuscript.

## Notes

Present address:

Quincy Bizjak, Patrick Halley: DNA Nanobots Inc., Powell, OH 43065, USA

