## Supplementary Data for "In‑Situ ssDNA Isolation from dsDNA Sources as a Streamlined Pathway to DNA Origami Assembly and Testing"

**Supporting data for Figure 1**

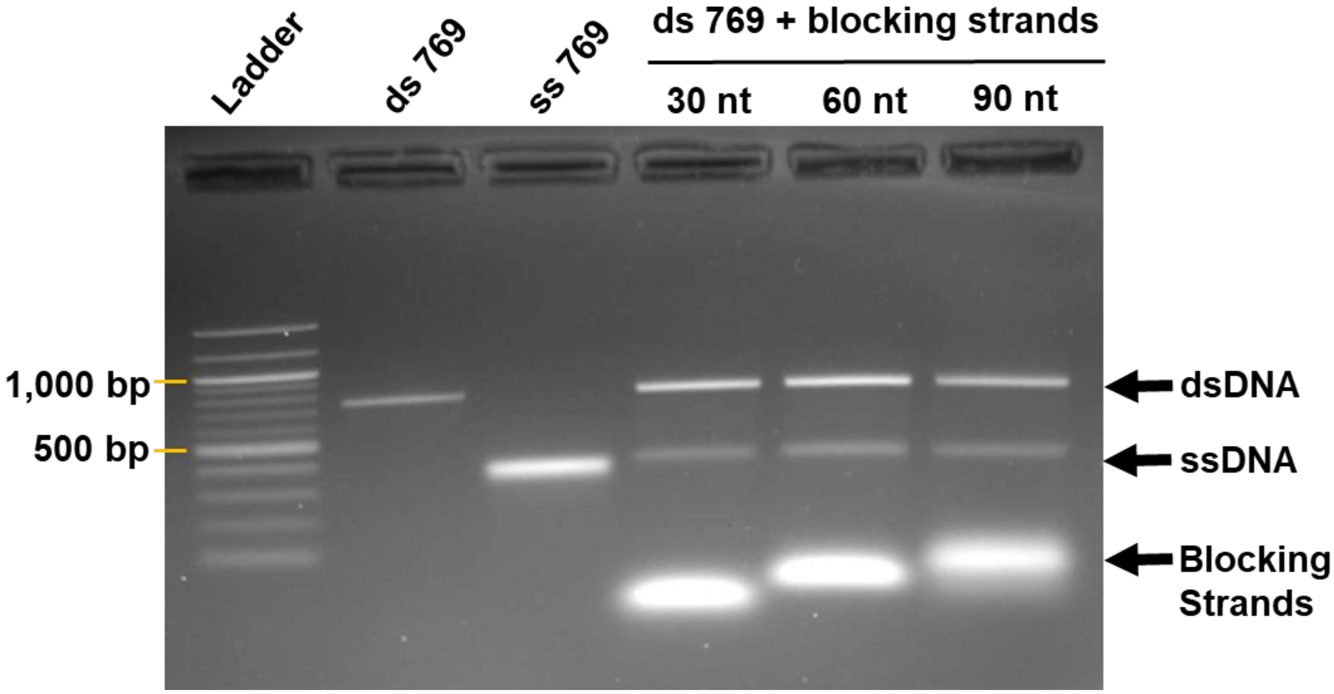


**Figure S1.** Agarose gel (2%) electropherogram (in 1X TBE with 11 mM $MgCl_2$) showing ssDNA release from a 769 bp dsDNA template using blocking strands of different lengths (30, 60, or 90 nt). Blocking strands were used in 10-fold excess over the dsDNA template. Samples were melted at **98°C** for **70 s**, followed by incubation at **64°C** for **5 min** after which the temperature was dropped to **20°C.** Samples were then evaluated by gel electrophoresis. Yield of ssDNA was slightly higher with 60 nt blocking strands compared to 30 nt blocking strands. Yield of ssDNA with 90 nt blocking strands was similar to 60 nt, but we moved forward with 60 nt strands since they have a lower cost.

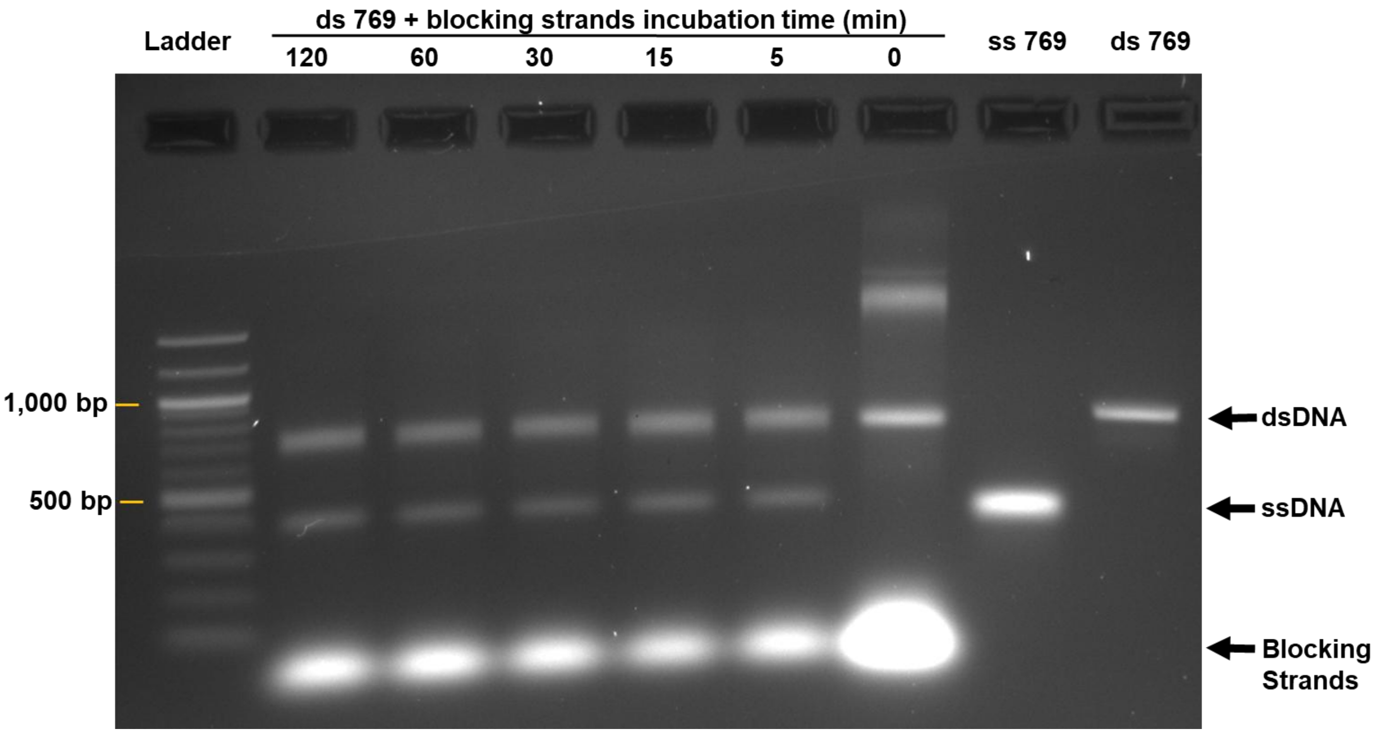


**Figure S2.** Agarose gel (2%) electropherogram (in 1X TBE with 11 mM $MgCl_2$) showing ssDNA release from a 769 bp dsDNA template using 10-fold excess **60 nt** blocking strands at different annealing times. The 0 min sample was heated to **98°C** for **70 s**, then kept at room temperature. The other samples were held at an annealing temperature of **64°C** for varying times prior to decreasing to room temperature. Results show 5 min is sufficient time for blocking strands to anneal and isolate the ssDNA.

**Sanger Sequencing (Figure 2).** The ssDNA generated in **Figure 2** (1-fold blocking strands, incubated for 5 min at 64/74/84°C) were analyzed via Sanger sequencing (Azenta) to verify homology with the ss2118 enzymatically prepared control strand. To isolate the ss2118 products, samples were gel-extracted (Freeze 'N Squeeze, BioRad) and concentrated with Ethanol precipitation. Representative trace chromatograms are shown below in Figure S3 and the primer sequences provided for Sanger sequencing are listed in **Table S5**.

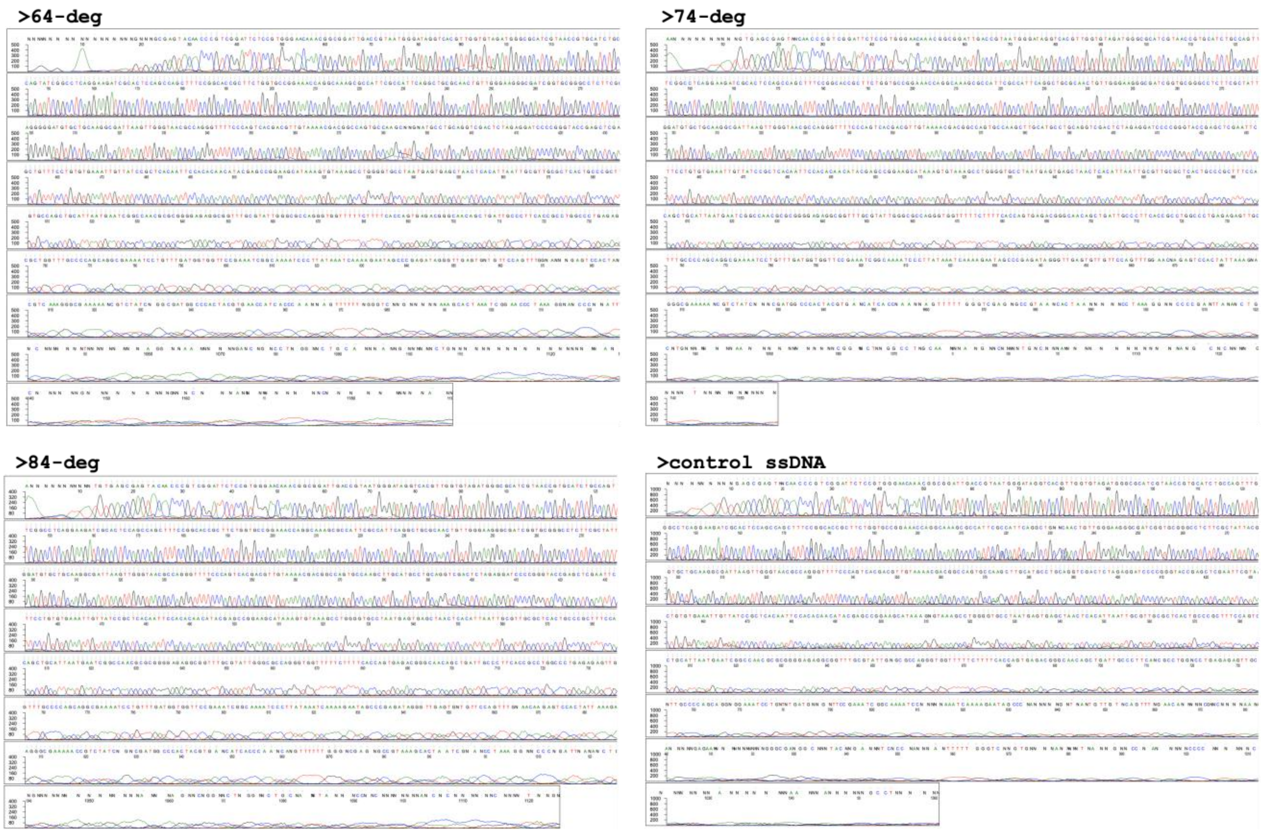


**Figure S3.** Trace chromatograms for released ss2118 following gel extraction. The reaction contained 1-fold blocking strands and underwent a 5 min incubation at 64, 74, or 84°C.

**>64deg-**
2118_primers_E05.ab1NNNNNNNNNNNNNNNNNNGNNNGCGAGTACAACCCGTCGGATTCTCCGTGGGAACAAACGGCGGATTGA
CCGTAATGGATAGGTCACGTTGGTGTAGATGGGCGCATCGTAACCGTGCATCTGCCAGTTTGAGGGGACGACGACAGTATCGGCCTCA
GGAAGATCGCACTCCAGCCAGCTTTCCGGCACCGCTTCTGGTGCCGGAAACCAGGCAAAGCGCCATTCGCCATTCAGGCTGCGCAACTG
TTGGGAAGGGCGATCGGTGCGGGCCTCTTCGCTATTACGCCAGCTGGCGAAAGGGGGATGTGCTGCAAGGCGATTAAGTTGGGTAACGC
CAGGGTTTTCCCAGTCACGACGTTGTAAAACGACGGCCAGTGCCAAGCNNGNATGCCTGCAGGTCGACTCTAGAGGATCCCCGGGTACC
GAGCTCGAATTCGTAATCATGGTCATAGCTGTTTCCTGTGTGAAATTGTTATCCGCTCACAATTCCACACAACATACGAGCCGGAAGCA
TAAAGTGTAAAGCCTGGGGTGCCTAATGAGTGAGCTAACTCACATTAATTGCGTTGCGCTCACTGCCCGCTTTCCAGTCGGGAAACCTG
TCGTGCCAGCTGCATTAATGAATCGGCCAACGCGCGGGGAGAGGCGGTTTGCGTATTGGGCGCCAGGGTGGTTTTTCTTTTCACCAGTG
AGACGGGCAACAGCTGATTGCCCTTCACCGCCTGGCCCTGAGAGAGTTGCAGCAAGCGGTCCACGCTGGTTTGCCCCAGCAGGCGAAAA
TCCTGTTTGATGGTGGTTCCGAAATCGGCAAAATCCCTTATAAATCAAAAGAATAGCCCGAGATAGGGTTGAGTGNTGTTCCAGTTTGG
NANNNGAGTCCACTANTAAAGAACGTGGACTCCAACGTCAAAGGGCGAAAAANCGTCTATCNGGCGATGGCCCACTACGTGAACCATCA
CCCAANNAGTTTTTTNGGGTCNNGNNNNNAAAGCACTAAATCGGAACCCTAAAGGNANCCCNNATTTNNNNCTGANGNNAANCNNNNNN
NTNNNNNNNNNNAGGNNAANNNNNNGANCNGNCCTNGGNNCTGCANNNANNGNNNNNNCTGNNNNNNNNNNNNNNNNNNNNANNNNNNNNN
NNCNNNNNNGNNNNNNNNNNGNNNCNNNANNNNNNNNNNNNNCNNNNNNNNNNNANNGN

**>74-deg-**
2118_primers_F05.ab1AANNNNNNNNNNGTGAGCGAGTNNCAACCCGTCGGATTCTCCGTGGGAACAAACGGCGGATTGACCGTA
ATGGGATAGGTCACGTTGGTGTAGATGGGCGCATCGTAACCGTGCATCTGCCAGTTTGAGGGGACGACGACAGTATCGGCCTCAGGAAG
ATCGCACTCCAGCCAGCTTTCCGGCACCGCTTCTGGTGCCGGAAACCAGGCAAAGCGCCATTCGCCATTCAGGCTGCGCAACTGTTGGG
AAGGGCGATCGGTGCGGGCCTCTTCGCTATTACGCCAGCTGGCGAAAGGGGGATGTGCTGCAAGGCGATTAAGTTGGGTAACGCCAGGG

TTTTCCCAGTCACGACGTTGTAAAACGACGGCCAGTGCCAAGCTTGCATGCCTGCAGGTCGACTCTAGAGGATCCCCGGGTACCGAGCT
CGAATTCGTAATCATGGTCATAGCTGTTTCCTGTGTGAAATTGTTATCCGCTCACAATTCCACACAACATACGAGCCGGAAGCATAAAG
TGTAAAGCCTGGGGTGCCTAATGAGTGAGCTAACTCACATTAATTGCGTTGCGCTCACTGCCCGCTTTCCAGTCGGGAAACCTGTCGTG
CCAGCTGCATTAATGAATCGGCCAACGCGCGGGGAGAGGCGGTTTGCGTATTGGGCGCCAGGGTGGTTTTTCTTTTCACCAGTGAGACG
GGCAACAGCTGATTGCCCTTCACCGCCTGGCCCTGAGAGAGTTGCAGCAAGCGGTCCACGCTGGTTTGCCCCAGCAGGCGAAAATCCTG
TTTGATGGTGGTTCCGAAATCGGCAAAATCCCTTATAAATCAAAAGAATAGCCCGAGATAGGGTTGAGTGTTGTTCCAGTTTGGAACNA
GAGTCCACTATTAAAGNACGTGGNNTCCAACGTCAAGGGCGAAAAANCGTCTATCNNNCGATGGCCCACTACGTGANCATCACCNAANN
AGTTTTTGGGTCGAGNGCCGTAANCACTAANNNNNCCTAAAGGNNCCCCGANTTANANCTGANGNNNANNNNNACNTGNNNNNNNNAAN
NNNNNNNNNNNCGGNNCTNNGGCCTNGCAANNNNANGNNCNNNNNTGNCNNNANNNNNNNNNNNNNNNANGCNCNNNNNGNNNNNACNAN
GNNNNTNNNNNNNNNNNNNNANN

**>84-deg-**
2118_primers_G05.ab1ANNNNNNNNNNNTGTGAGCGAGTACAACCCGTCGGATTCTCCGTGGGAACAAACGGCGGATTGACCGTA
ATGGGATAGGTCACGTTGGTGTAGATGGGCGCATCGTAACCGTGCATCTGCCAGTTTGAGGGGACGACGACAGTATCGGCCTCAGGAAG
ATCGCACTCCAGCCAGCTTTCCGGCACCGCTTCTGGTGCCGGAAACCAGGCAAAGCGCCATTCGCCATTCAGGCTGCGCAACTGTTGGG
AAGGGCGATCGGTGCGGGCCTCTTCGCTATTACGCCAGCTGGCGAAAGGGGGATGTGCTGCAAGGCGATTAAGTTGGGTAACGCCAGGG
TTTTCCCAGTCACGACGTTGTAAAACGACGGCCAGTGCCAAGCTTGCATGCCTGCAGGTCGACTCTAGAGGATCCCCGGGTACCGAGCT
CGAATTCGTAATCATGGTCATAGCTGTTTCCTGTGTGAAATTGTTATCCGCTCACAATTCCACACAACATACGAGCCGGAAGCATAAAG
TGTAAAGCCTGGGGTGCCTAATGAGTGAGCTAACTCACATTAATTGCGTTGCGCTCACTGCCCGCTTTCCAGTCGGGAAACCTGTCGTG
CCAGCTGCATTAATGAATCGGCCAACGCGCGGGGAGAGGCGGTTTGCGTATTGGGCGCCAGGGTGGTTTTTCTTTTCACCAGTGAGACG
GGCAACAGCTGATTGCCCTTCACCGCCTGGCCCTGAGAGAGTTGCAGCAAGCGGTCCACGCTGGTTTGCCCCAGCAGGCGAAAATCCTG
TTTGATGGTGGTTCCGAAATCGGCAAAATCCCTTATAAATCAAAAGAATAGCCCGAGATAGGGTTGAGTGNTGTTCCAGTTTGNAACAA
GAGTCCACTATTAAAGAACGTGGNNTCCNACGTCAAAGGGCGAAAAACCGTCTATCNGNCGATGGCCCACTACGTGANCATCACCCAAN
CANGTTTTTTGGGNCGAGNGCCGTAAAGCACTAATCGNANCCTAAAGGNNCCCNGATTNANANCTGACGGGGAANNNNNANGNNNNNNN
NNNNNNNNANNNAGNNCNGGNNCTNGGNNCTGCNANNTANNNCCNNCNNNNNNNNNANCNCNNNNNNCNNNNTNNGNNN

**>ss2118 control-**
2118_primers_H05.ab1NNNNNNNNNNGAGCGAGTNNCAACCCGTCGGATTCTCCGTGGGAACAAACGGCGGATTGACCGTAATGG
GATAGGTCACGTTGGTGTAGATGGGCGCATCGTAACCGTGCATCTGCCAGTTTGAGGGGACGACGACAGTATCGGCCTCAGGAAGATCG
CACTCCAGCCAGCTTTCCGGCACCGCTTCTGGTGCCGGAAACCAGGCAAAGCGCCATTCGCCATTCAGGCTGNNCAACTGTTGGGAAGG
GCGATCGGTGCGGGCCTCTTCGCTATTACGCCAGCTGGCGAAAGGGGGATGTGCTGCAAGGCGATTAAGTTGGGTAACGCCAGGGTTTT
CCCAGTCACGACGTTGTAAAACGACGGCCAGTGCCAAGCTTGCATGCCTGCAGGTCGACTCTAGAGGATCCCCGGGTACCGAGCTCGAA
TTCGTAATCATGGTCATAGCTGTTTCCTGTGTGAAATTGTTATCCGCTCACAATTCCACACAACATACGAGCCGGAAGCATAAAGNGTA
AAGCCTGGGGTGCCTAATGAGTGAGCTAACTCACATTAATTGCGTTGCGCTCACTGCCCGCTTTCCAGTCGGGGANNCCTGTCGTGCCA
GCTGCATTAATGAATCGGCCAACGCGCGGGGAGAGGCGGTTTGCGTATTGNGCGCCAGGGTGGTTTTTCTTTTCACCAGTGAGACGGGC
AACAGCTGATTGCCCTTCANCGCCTGGNCCTGAGAGAGTTGCAGCAAGCGGTCCANGCTGGNTTGCCCCAGCAGGNGGAAATCCTGNTN
TGATGNNGNTTCCGAAATCGGCAAAATCCNNNNNAAATCAAAAGAATAGCCCNANNNNNGNTNANTGTTGTNCAGTTTNGAACANNNNN
CGNNCNNNNAANAACNNGNNTCCANGTCNANNNNNGAGAANNNNNNNNNNNNNGGGCGANGGCNNNTACNNGANNNTCNCCNANNNANT
TTTTGGGTCNNGTGNNNNANNNNNTNANNGNNCCNANNNNNCCCCNNNNNNCTNNNNGNNNNNNNNNNNNNANNNNNNNNNAANNNANNN
NNNGCCTNNNNNNGNNN

**Table S5: Sequencing results for Figure S4.**

| Sample | Quality Score+ | Contiguous Read Length |
|---|---|---|
| 64°C – 2118 | 37 | 851 |
| 74°C – 2118 | 40 | 870 |
| 84°C – 2118 | 40 | 961 |
| Control ssDNA | 29 | 763 |

+Quality Score ≥ 40 and Contiguous Read Length > 500 = Good trace results
Quality Score 25-39 and Contiguous Read Length > 500 = May have a high background signal, but the trace may still have usable sequence present.

**Quantification with ImageJ.** Gel images were imported as .jpeg files without prior contrast enhancement. In cases where there were multiple bands per lane, intensity profiles were defined manually as shown in Figure S4.

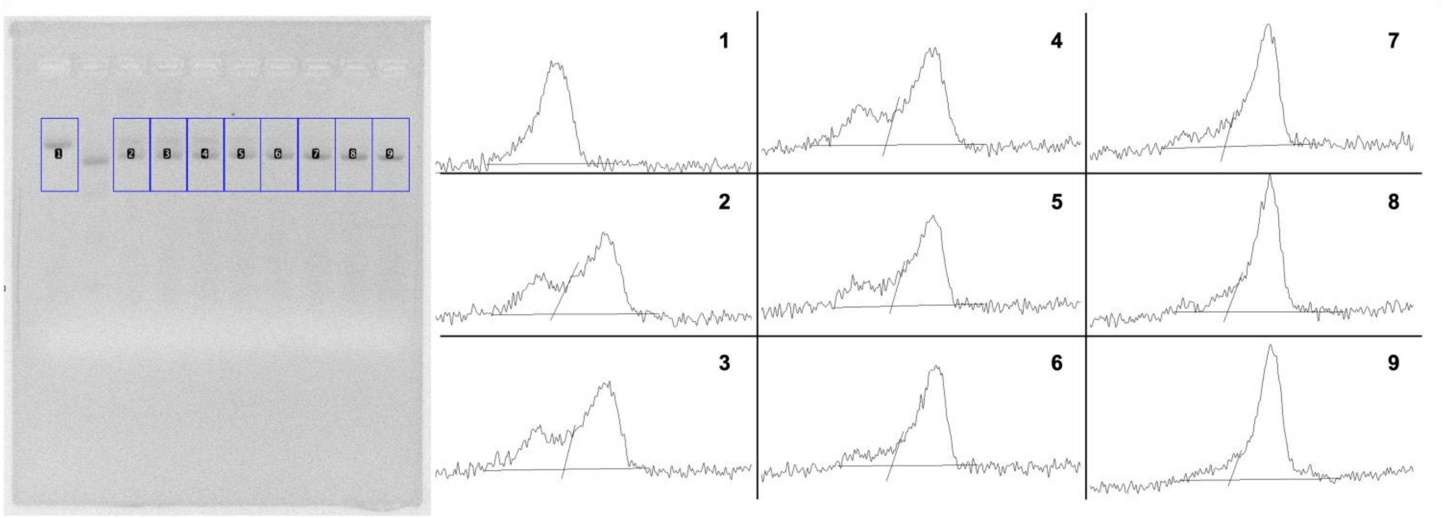


**Figure S4.** Representative example for band intensity quantification used to calculate released ssDNA yields. The gel shown corresponds to a 74°C, 5 min incubation of the 2,118 dsDNA template.

**Table S6:** Experimental specifics for ssDNA release corresponding to each DO.

| DO | dsDNA template concentration (nM) | Blocking strand excess | Annealing temperature (°C) | Incubation time (min) |
|---|---|---|---|---|
| **2,118-monolayer** | 10 | 10 | 64 | 5 |
| **phiX-nanotube** | 10 | 10 | 64 | 5 |
| **pUC19-monolayer** | 8 | 25 | 64 | 5 |
| **Lambda 4,674** | 10 | 20 | 64 | 5 |
| **Lambda 4,845** | 10 | 20 | 64 | 5 |
| **Lambda 5,582** | 10 | 20 | 64 | 5 |
| **Lambda 7,088** | 10 | 20 | 64 | 5 |
| **Lambda 8,013** | 10 | 20 | 64 | 5 |
| **Lambda 15,101** | 1.2 | 20 | 64 | 5 |
| **eGFP-DO** | 10 | 5 | 58 | 1 |

**Supporting data for Figure 3**

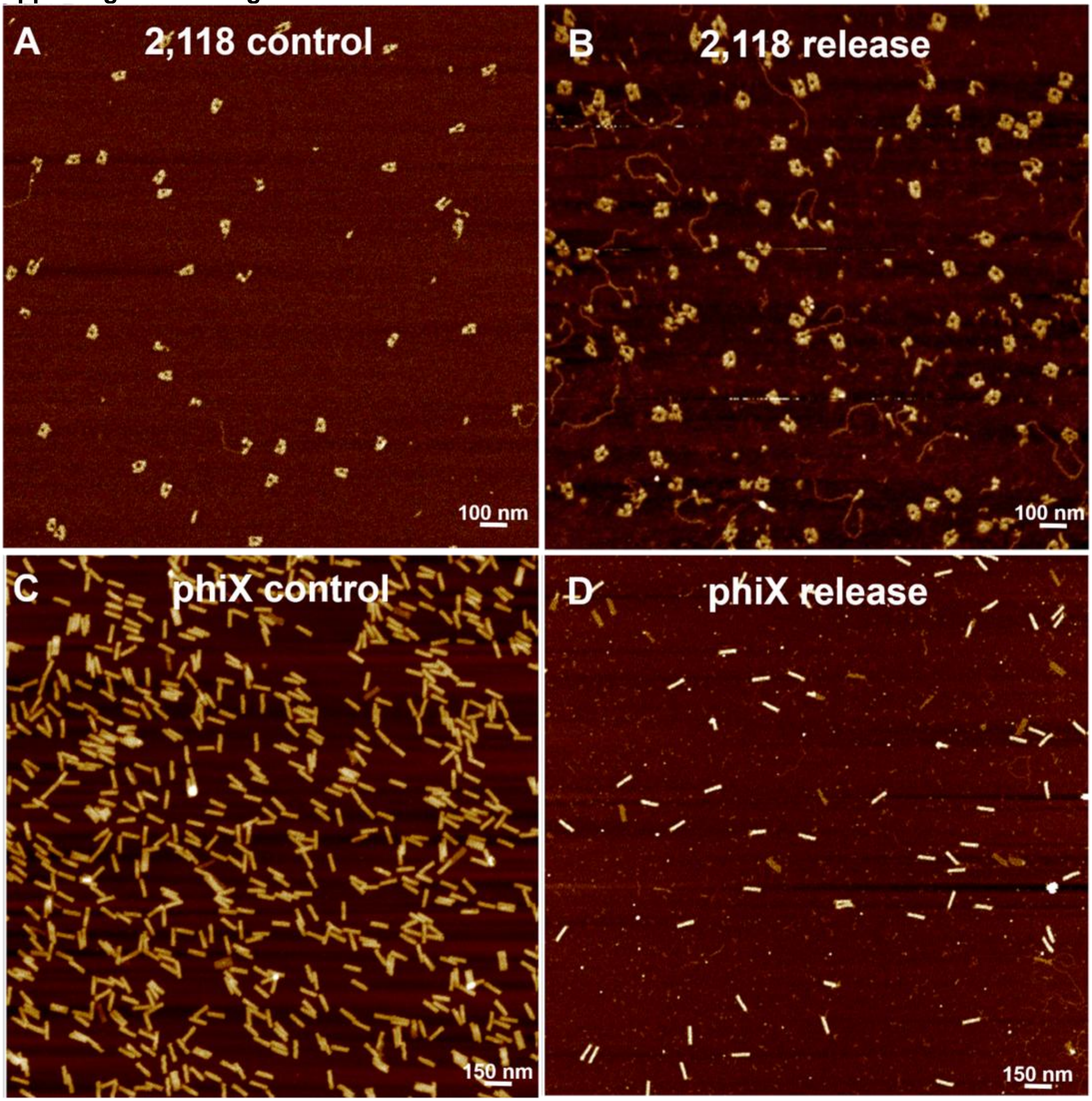


**Figure S5.** Representative AFM images corresponding to results in Figure 3. (A) Gel purified 2,118 monolayer DOs folded with enzymatically prepared ssDNA. (B) 2,118 monolayer DOs folded from released ssDNA. (C) PEG purified phiX-nanotube folded from control ssDNA plasmid. (D) DO nanotube folded with ssDNA released from the dsDNA precursor.

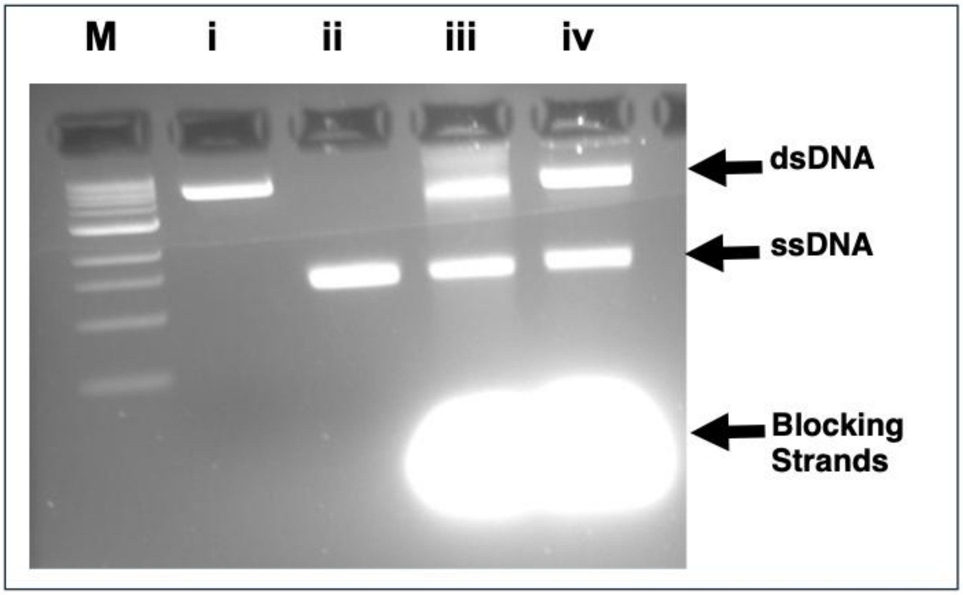


**Figure S6.** Agarose gel (2%) electropherogram showing ssDNA release and purification from phiX174 dsDNA template, used for DO rod assembly in Figure 3F. Lanes are as follows- M: molecular weight ladder; i: phiX174 linearized dsDNA template; ii: phiX174 ssDNA control; iii: ssDNA released from 10-fold blocking strands using purified linear phiX174 dsDNA template; iv: ssDNA released in the same manner but using unpurified linear phiX174 dsDNA template.

**Supporting data for Figure 4**

| | | | | | | |
|---|---|---|---|---|---|---|
| + | + | + | + | + | | ds2118 |
| | | | | | + | ss2118 |
| | + | + | + | + | | Blocking strands |
| + | + | + | + | + | + | Staples |
| 5 | 5 | 10 | 5 | 10 | 10 | $Mg^{2+}$ (mM) |
| | 1:1:10 | 1:1:10 | 1:3:10 | 1:3:10 | | |

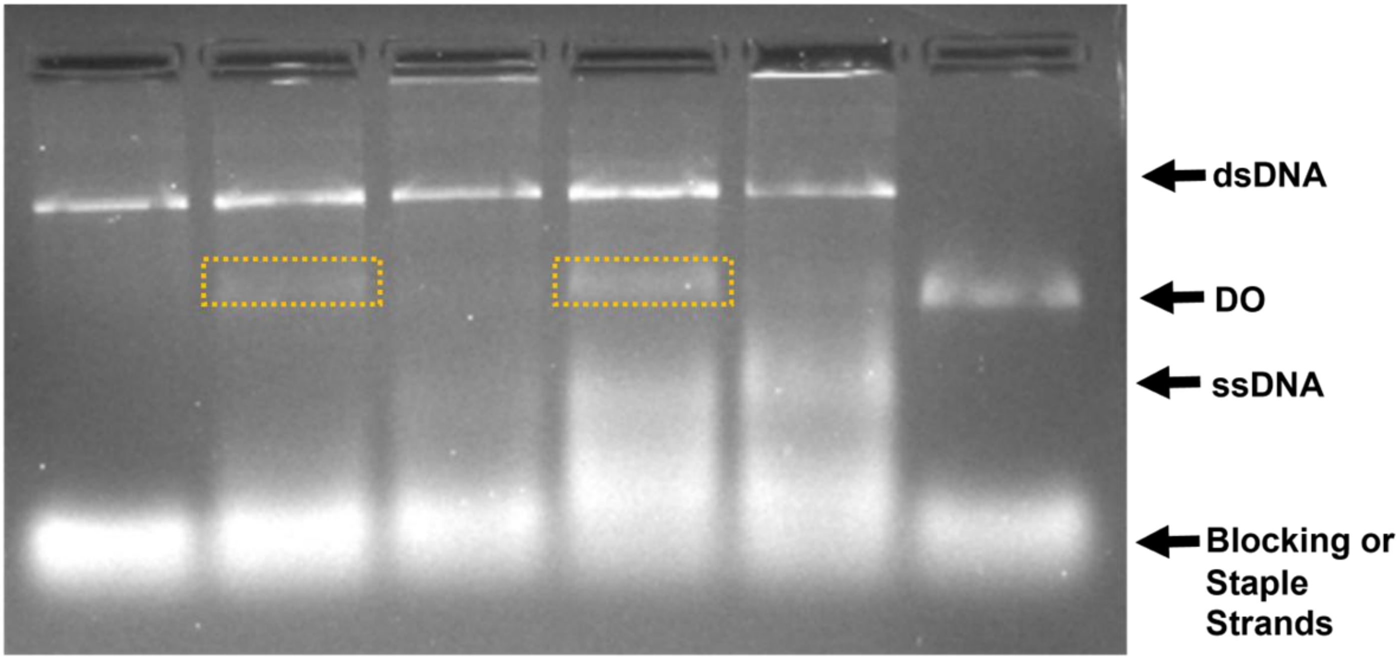


**Figure S7.** Agarose gel (2%) electropherogram showing one-pot assembly of 2,118 based monolayer DO at different blocking strand concentrations (ratio = template : blocking strands : staple strands).

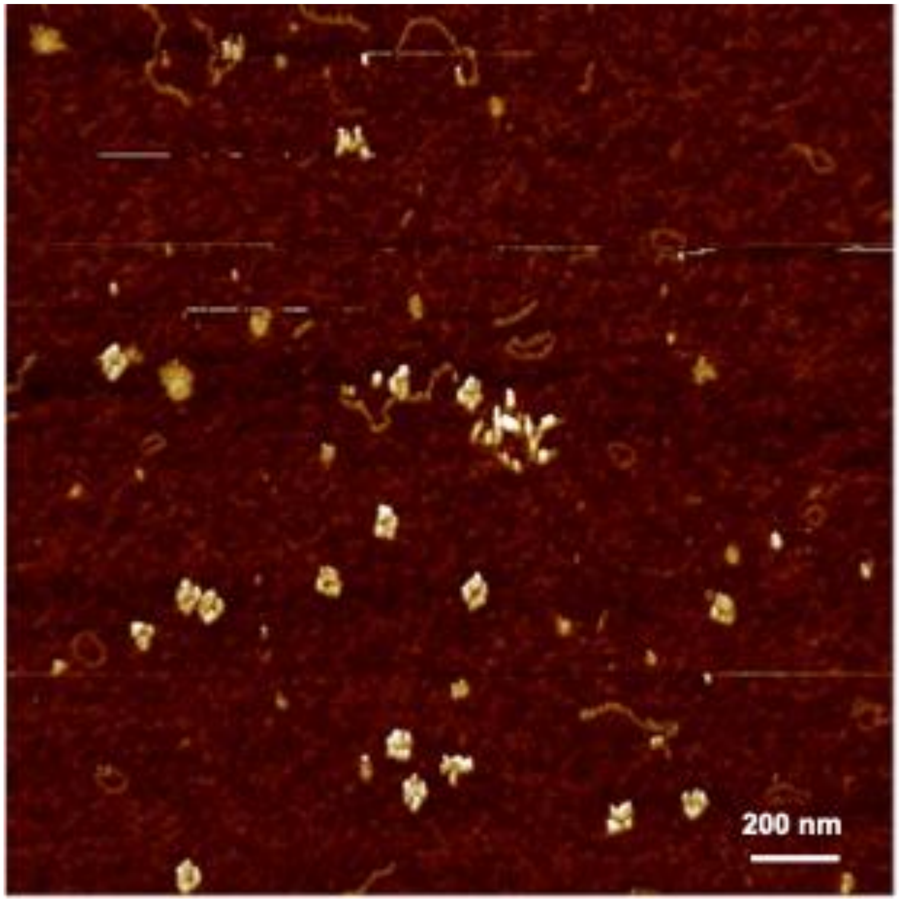


**Figure S8.** Representative AFM image corresponding to results shown in Figure 4B. Gel purified DOs generated through the one pot assembly protocol for 2,118 bp based monolayer DO.

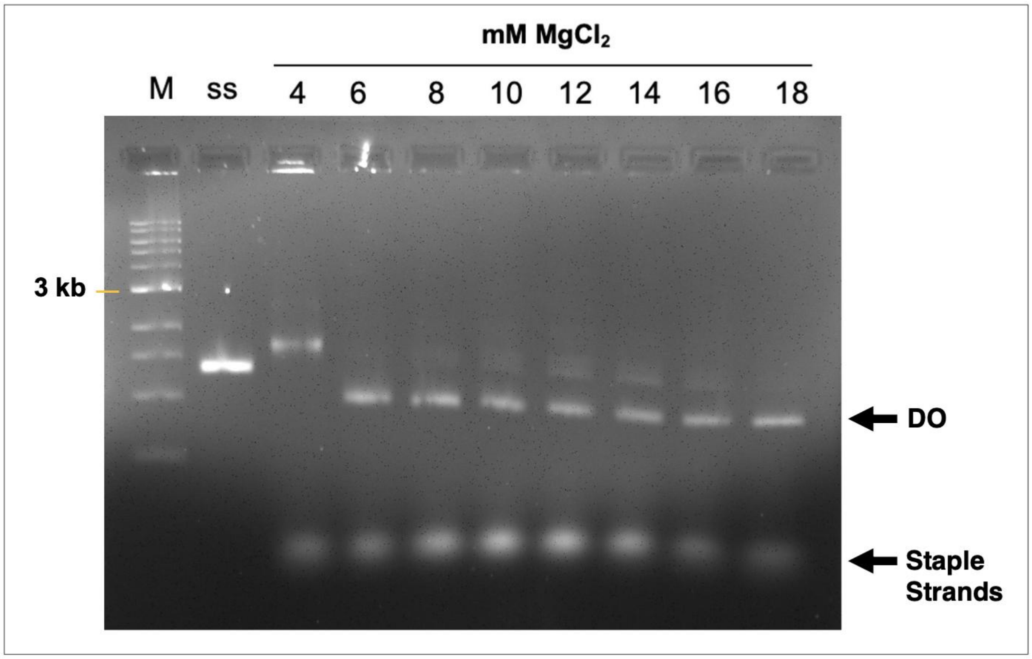

**Figure S9.** Prior to single-pot folding, we screened different $Mg^{2+}$ concentrations required for the one-pot assembly of phiX174 DO nanotube using 1% agarose gel electrophoresis.

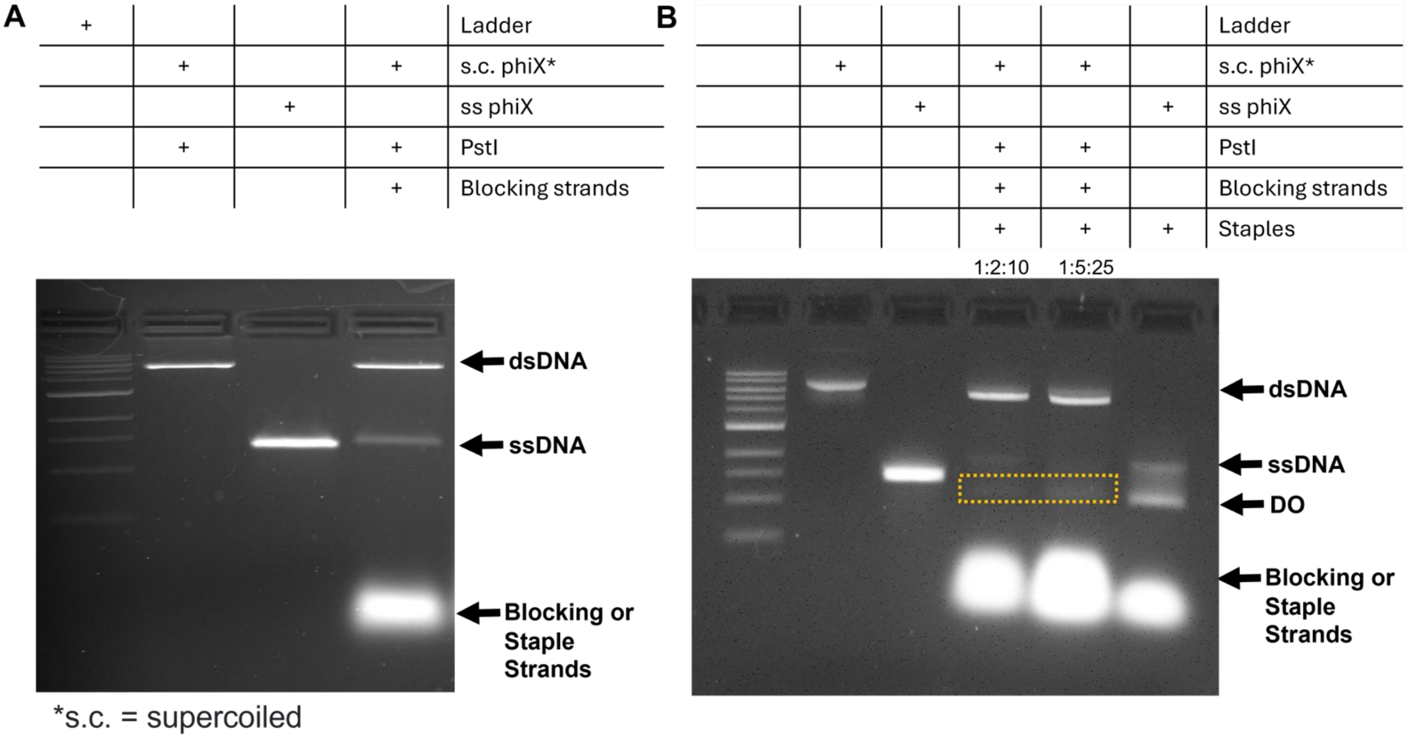


**Figure S10.** Agarose gel electropherograms showing one-pot assembly corresponding to the results in Figure 4C. (A) Agarose gel (2%, 1X TBE with 11 mM $MgCl_2$) showing phiX174 supercoiled (s.c.) dsDNA linearized, then a ssDNA control, followed by the release reaction using 3-fold excess blocking strands. (B) Agarose gel (1%, 1X TBE with 6 mM $MgCl_2$) showing one pot assembly of DO at different blocking strand concentrations (ratio = template : blocking strands : staple strands). The boxed bands were excised for AFM and TEM imaging.

| | | | | |
|---|---|---|---|---|
| + | | | | Ladder |
| | + | + | + | pUC19 |
| | + | + | + | PstI |
| | | + | + | Blocking strands |
| | | | + | Staples |

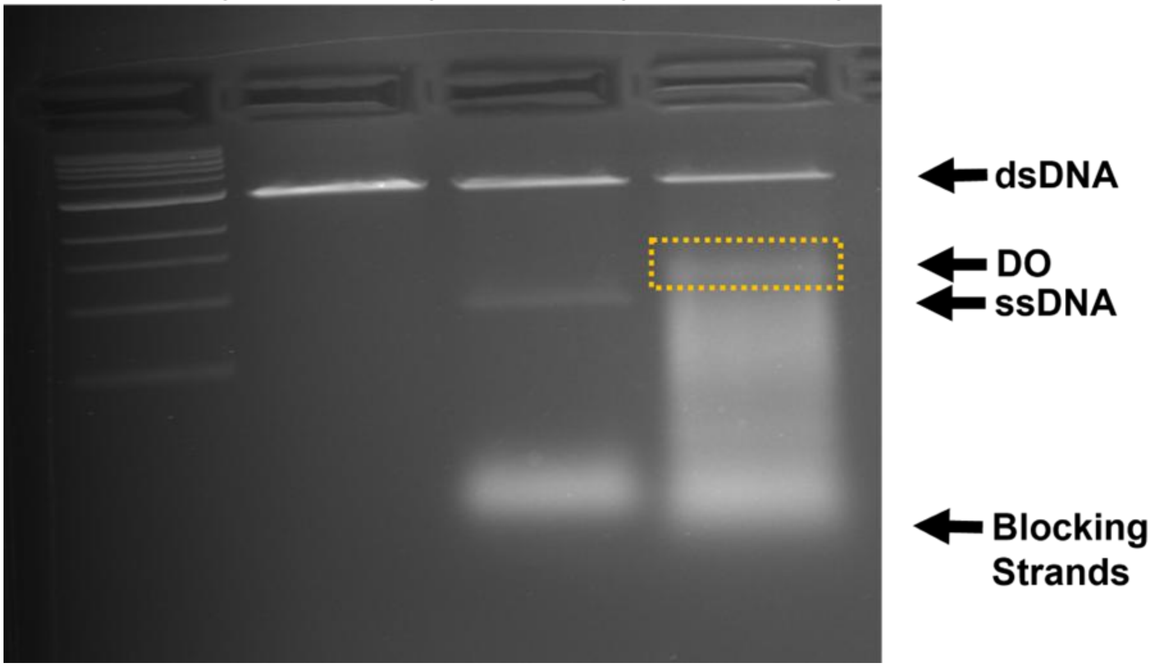


**Figure S11.** Agarose gel (2%) electropherogram showing one-pot assembly of pUC19 based monolayer DO. The boxed band was excised for AFM.

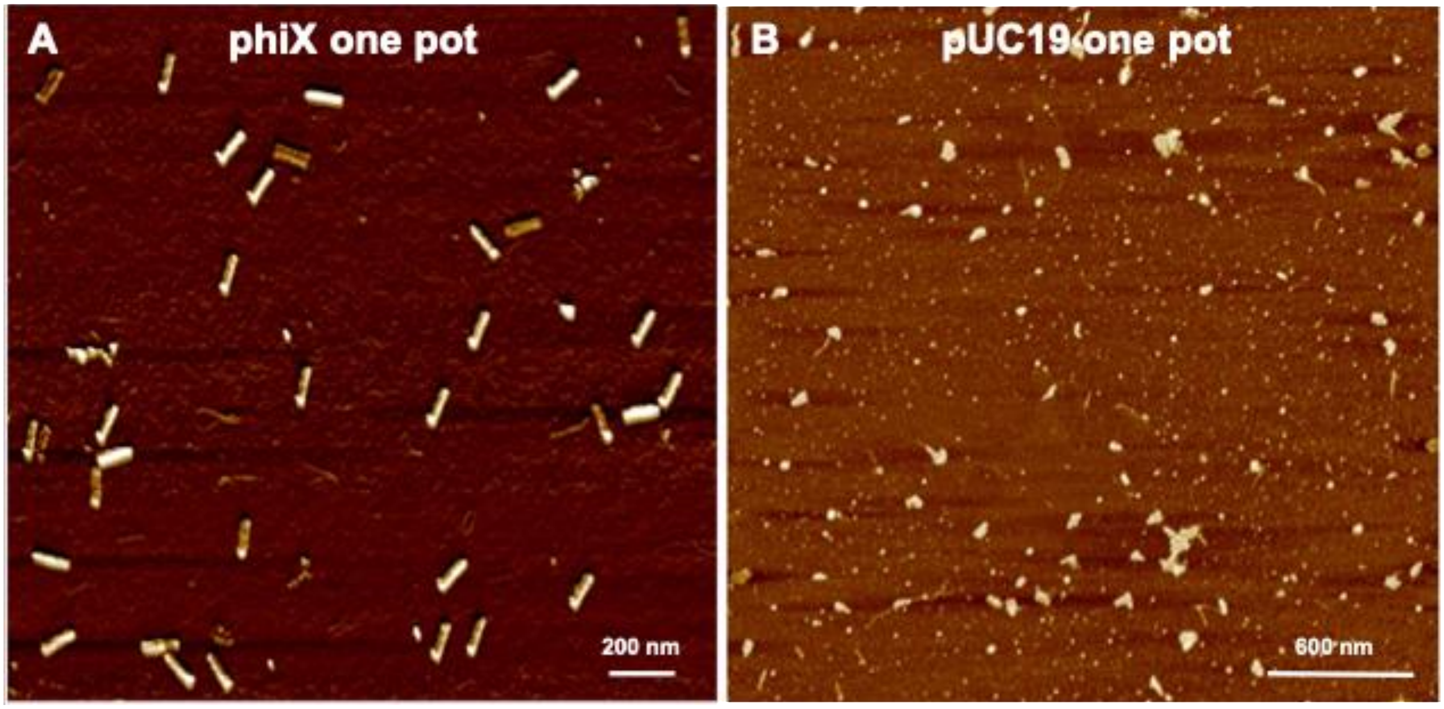

**Figure S12.** Representative AFM images corresponding to results shown in Figure 4C-D. Gel purified DOs generated through the one pot assembly protocol. (A) phiX rod, and (B) pUC19 monolayer.

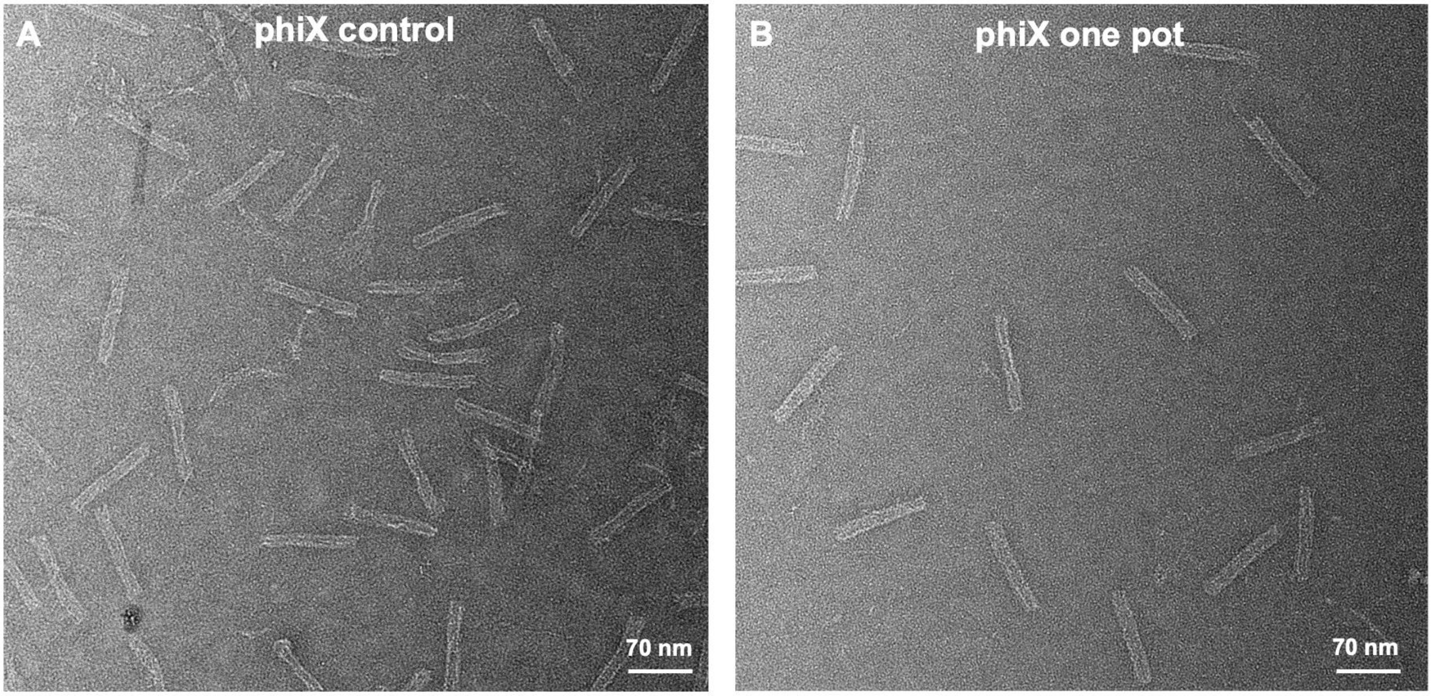


**Figure S13.** Representative TEM images of DO nanotube assembled using (A) control ssDNA scaffold, or (B) one-pot assembly.

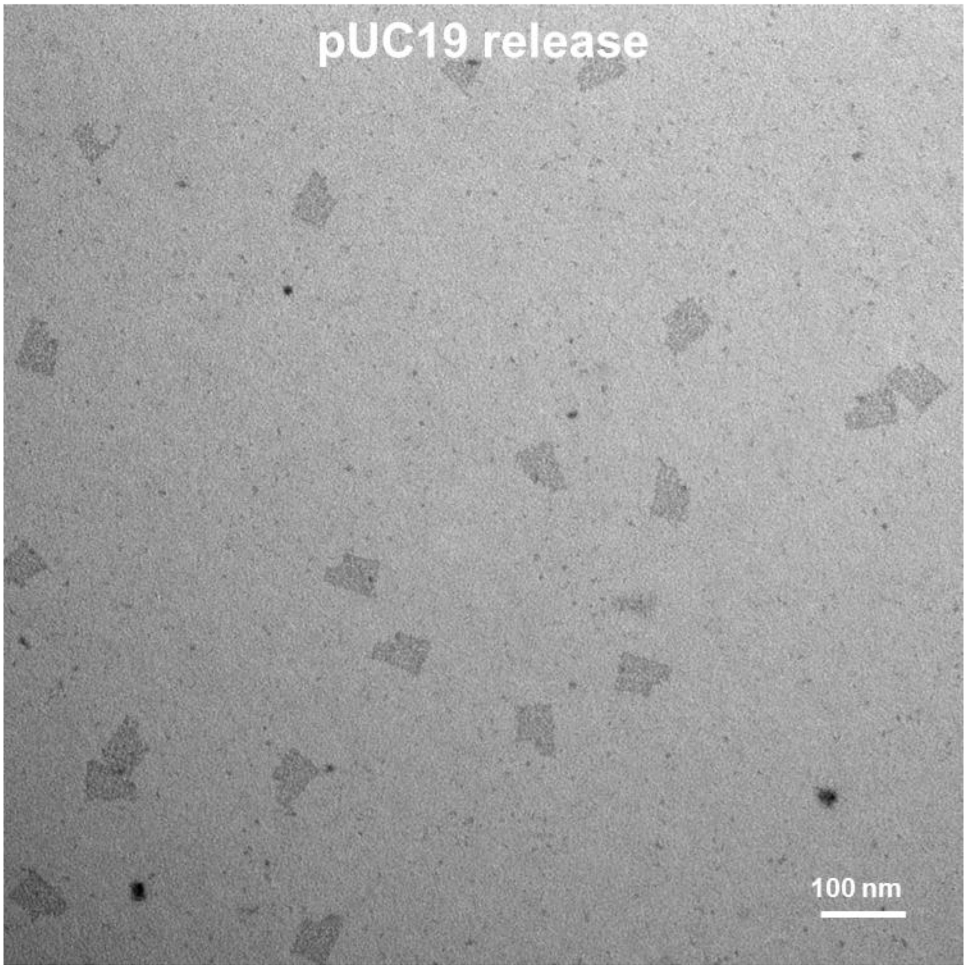

**Figure S14.** Representative TEM image of asymmetric DO rectangle assembled from released ssDNA.

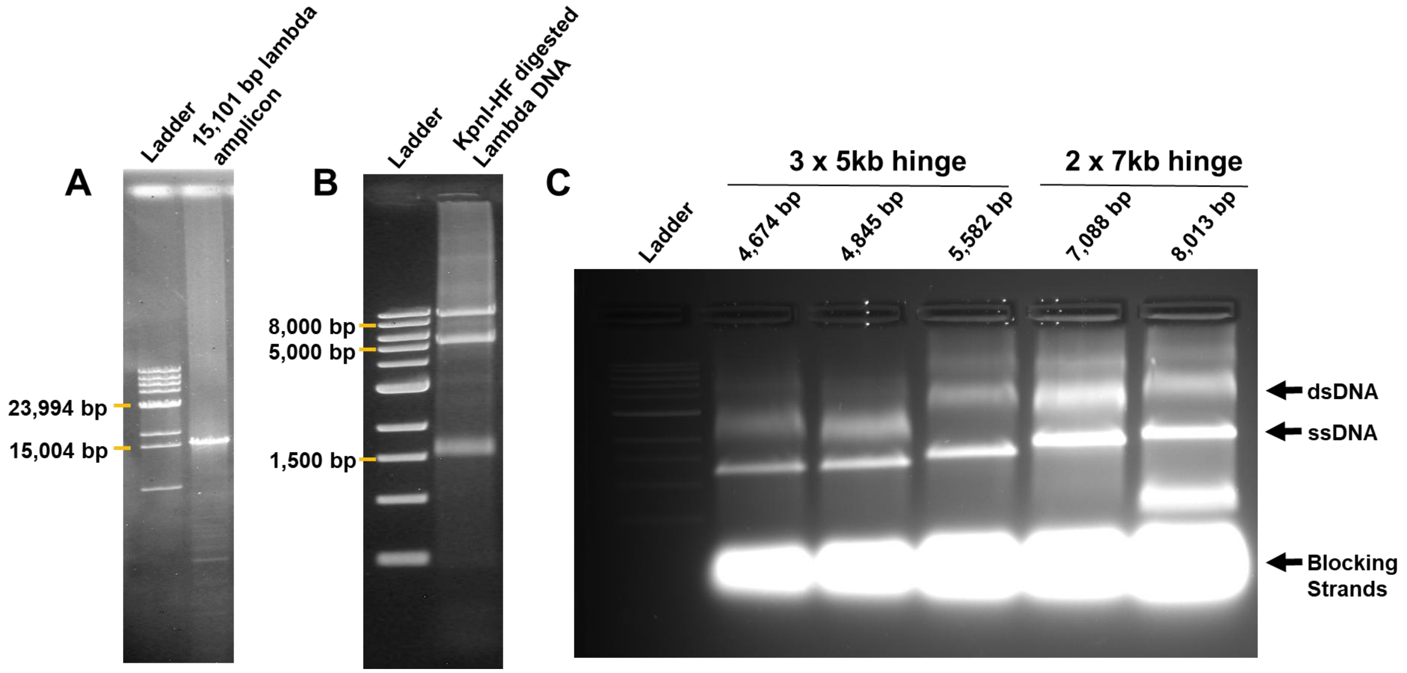


**Figure S15.** Agarose gel electropherogram for characterization of ssDNA release for hinge DO shown in Figure 5. (A) 15,101 bp PCR amplicon from lambda DNA. (B) KpnI digestion (cutting site: GGTAC|C) of the target amplicon showing the expected digestion pattern. Fragments are 8,587 bps, 5,011 bps, and 1,503 bps in size. C) AGE (1%) in 1X TBE with 11 mM $MgCl_2$ run for 1.5 h at 90 V. Template DNA (10 nM) was combined with 200 nM blocking strands and ssDNA release was facilitated by a 5 mi incubation at 64°C.

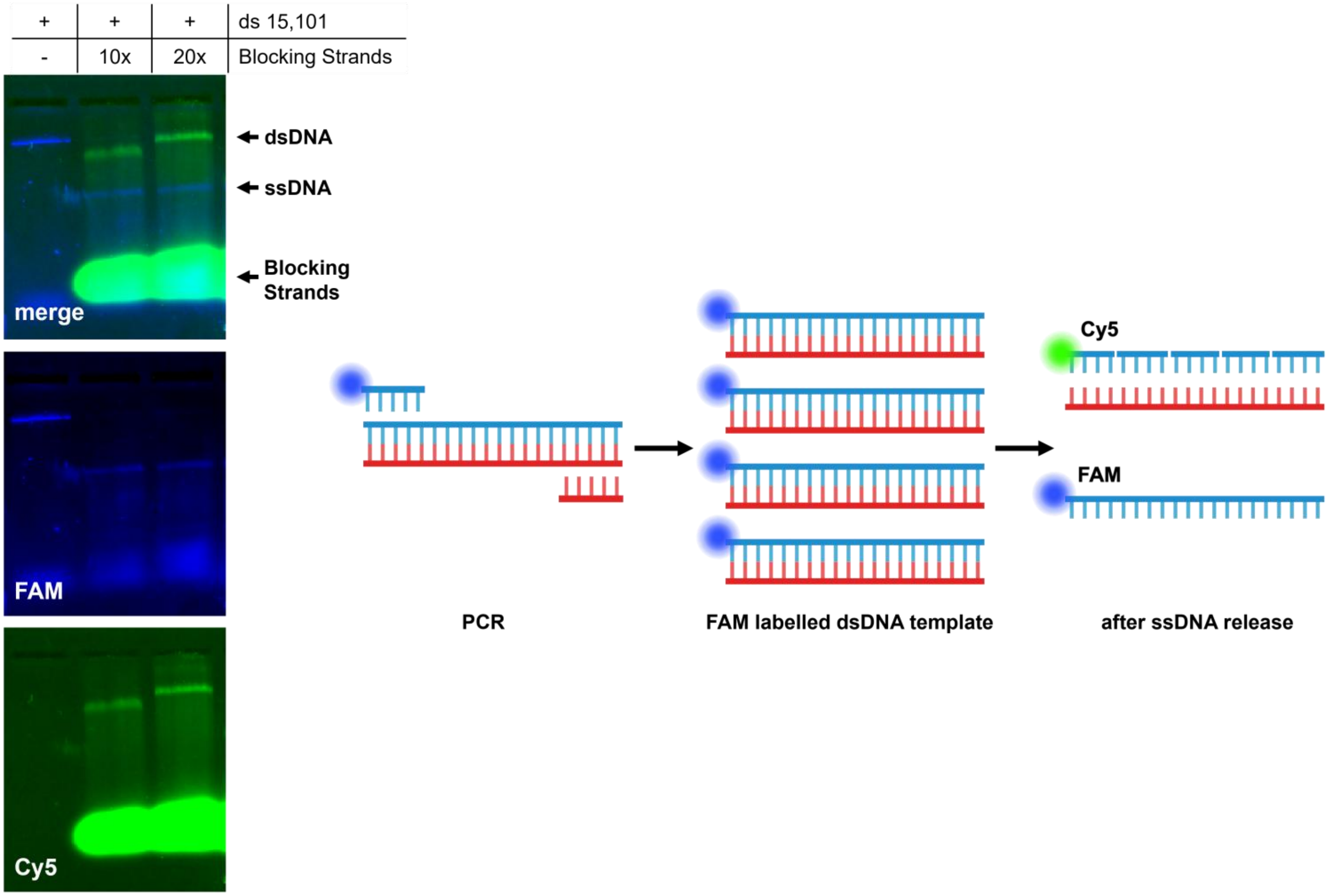


**Figure S16.** Agarose gel electropherogram for characterization of ssDNA release for hinge DO shown in Figure 5D. dsDNA template has been labelled with a FAM fluorophore and one of the blocking strands has been labelled with a Cy5 fluorophore to visualize migration of each species in the gel.

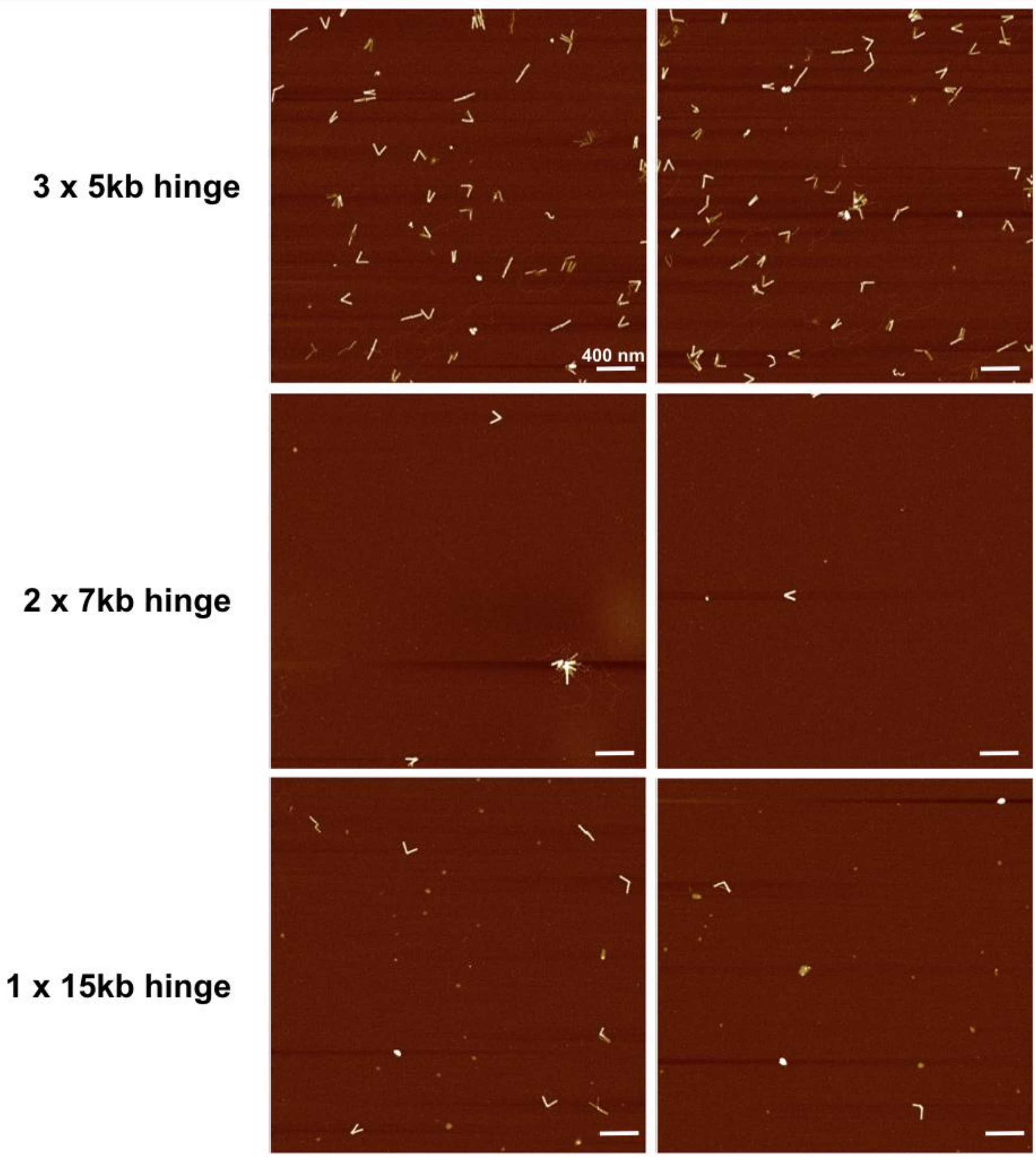


**Figure S17:** Representative AFM images of multi-scaffolded and large-scaffolded hinge DOs assembled from released ssDNA and shown in Figure 5. DOs were assembled with 0.6 nM scaffold, 20 nM staples, [12 mM $MgCl_2$], annealed with a 14 h thermal ramp, and were purified and concentrated via PEG precipitation. Scale bars = 400 nm.

**Supporting data for Figure 6**
**eGFP-encoding DO ssDNA scaffold release and subsequent purification**
Blocking strands were designed to be complementary to the coding/sense strand of the pCMV-T7-eGFP plasmid; as a result, the template/antisense strand was used as the scaffold for the resulting eGFP-DO. The oligos were designed to be 60 nt in length, and three of the blocking strands were biotinylated (**Table S3**) – one at each of the 3' and 5' ends of the template DNA, and one in the middle of the template DNA sequence.

Linear plasmid (10 nM), blocking strands (50 nM), $MgCl_2$ (5 mM), and TBE (1X) were combined in nuclease-free water; ssDNA was released by denaturation at 95°C for 10 min followed by an incubation at 58°C for 1 min.

**Streptavidin Magnetic Beads Preparation**. A wash buffer, recommended by published works, was prepared to rinse the streptavidin-coated magnetic beads just before use.[1, 2] Wash buffer comprised of 10 mM Tris HCl, 1 mM EDTA, 12 mM $MgCl_2$, 5 mM NaCl, 0.05% Tween20, and Nuclease-Free HyPure Molecular Biology Grade Water. To wash the beads, 500 µL wash buffer was combined with the streptavidin magnetic beads in a 1.5 mL Eppendorf vial, mixed by inverting the tube multiple times, and separated to discard the wash by placing it in a Cytiva MagRack 6 for 30 s. This wash step was repeated 6 times based on previously published work.[2]

The washed beads were then used for the removal of the dsDNA after the ssDNA release reaction and for subsequent 36hb purification following a previously optimized protocol. [2] Based on a functional assay by the vendor, the biotin binding capacity (BBC) of the beads is estimated to be 490-750 pmol/mg beads. We performed our calculations taking the BBC to be 500 pmol/mg. The purification procedure is as follows (illustrated in Figure S18):

**Removal of dsDNA from crude blocking solution.** Three blocking strands were biotinylated allowing for purification of the desired scaffold strand (dsDNA pull down).

1. 30-fold pmol excess bead BBC relative to pmol biotinylated blocking strands was combined with the crude blocking solution (Figure S18, lane iv). Generally, the volume of beads added to the crude blocking solution was 1 equivalent (v/v).
2. The mixture was placed on a Disruptor Genie set at 1000 rpm, for 3 h in a cold room (4°C).
3. Magnetic beads precipitation was used to separate the beads+captured dsDNA from the supernatant containing the released ssDNA scaffold. (Figure S18, lane v).

**Pull-down and photocleaving of 36hb**

1. 50-fold pmol excess bead BBC relative to pmol PC-biotin staple strands was combined with the annealed crude 36hb. Generally, the volume of beads added to the crude DO solution was 0.5 equivalents (v/v).
2. The bead+DNA NP mixture was placed on a Disruptor Genie set at 1000 rpm, for 3 h in a cold room (4°C).
3. Magnetic beads precipitation was used to separate the beads+captured DOs from the supernatant containing excess staple strands or other impurities by placing the vial in a MagRack for 30 s. Supernatant was removed and reserved in another vial for analysis (Figure S18, lane viii).
4. Beads with captured DOs were reconstituted in target buffer (1X TBE, 20 mM $MgCl_2$) at a 1:1 (v/v) ratio with the original (crude) DO volume.
5. To cleave the eGFP-DO from the beads, the sample was placed on a Nutating Mixer in a tube rack comprising 365 nm realUV LED Strip Lights for 60 m. The supernatant (containing pure

DO) was collected using magnetic bead precipitation for 30 s and the beads were subsequently discarded (Figure S18, lane vii).

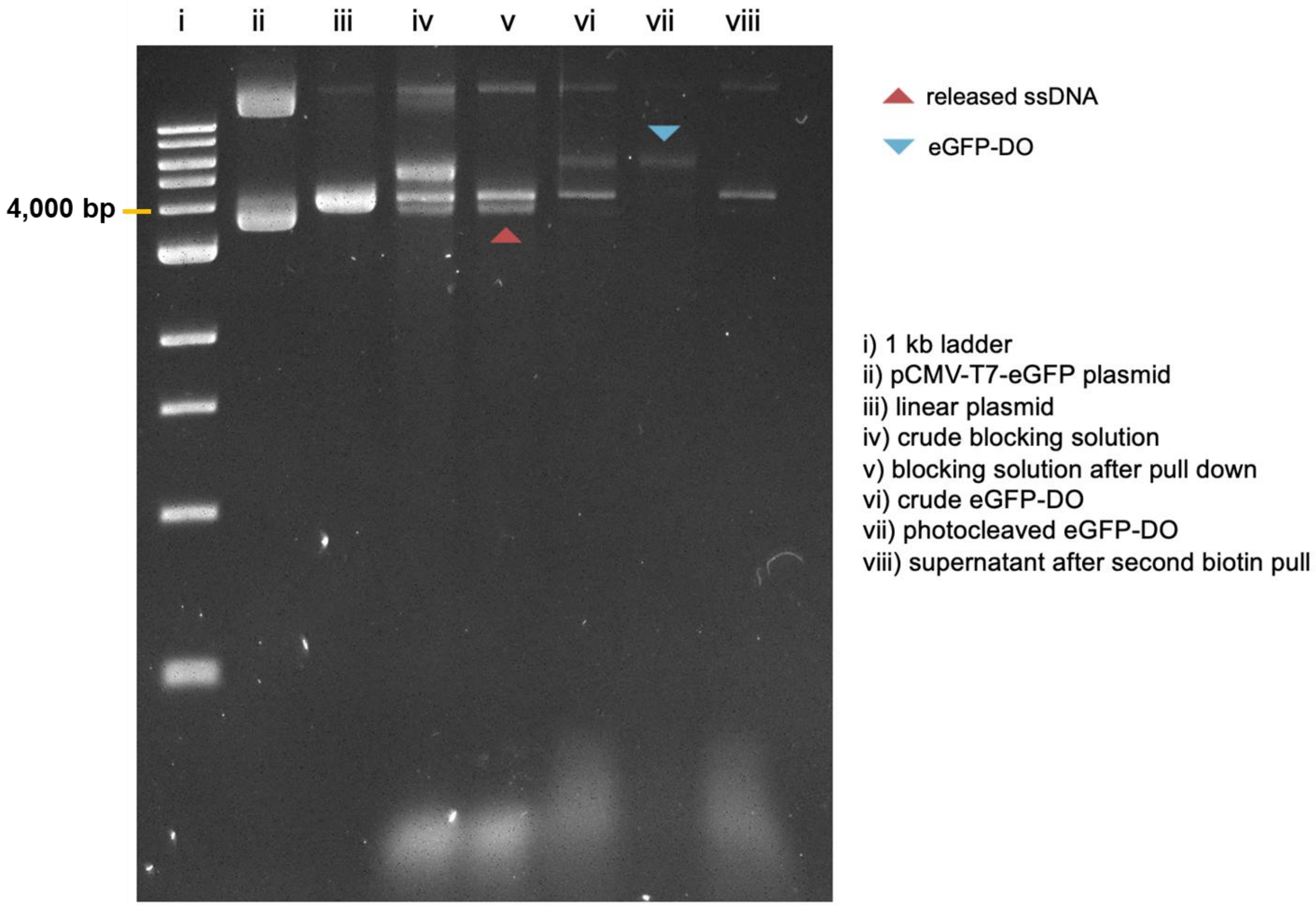


**Figure S18:** AGE characterization of eGFP-DO assembly and purification. Agarose gel (1%) ran at 90 V for 2 h in 1X TBE.

### GFP CHARACTERIZATION AND QUANTIFICATION

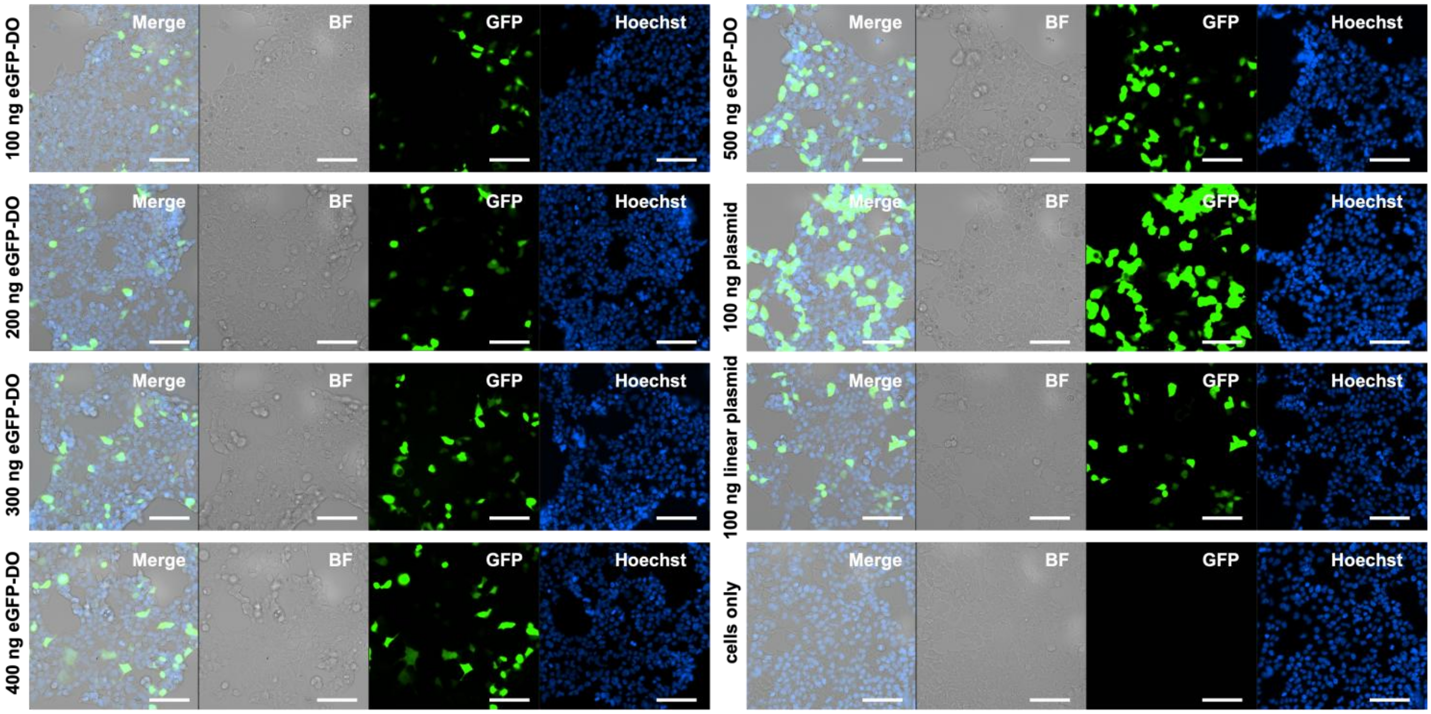


**Figure S19:** *In vitro* characterization of transfected eGFP-DO. HEK293 cells were transfected (Lipofectamine3000) with varying amounts of DO and were compared against supercoiled plasmid and linearized plasmid controls. Images were acquired 24 h post-transfection. Brightfield (BF) images were taken to assess cell morphology. eGFP expression is shown in the green channel (excitation 475 nm, emission 520 nm, laser power: 7), and the cell nuclei are visible in the Hoechst channel (excitation 385 nm, emission 461 nm, laser power: 7). Scale bar = 100 μm.

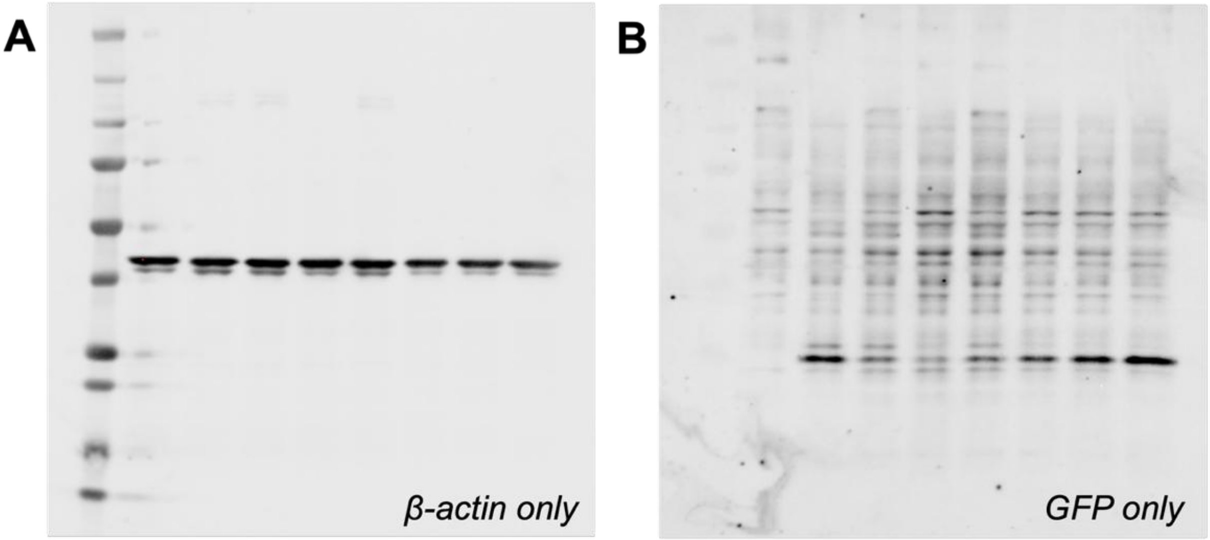


**Figure S20:** Raw Images of Western Blot of Figure 6D. The same membrane was imaged in two different channels. Sample order (left to right): Precision Plus Protein Ladder (BioRad, cat. #1610373), cells only, 100 ng pCMV-T7-eGFP plasmid, 100 ng linear pCMV-T7-eGFP plasmid, 100 ng eGFP-DO, 200 ng eGFP-DO, 300 ng eGFP-DO, 400 ng eGFP-DO, and 500 ng eGFP-DO. (A) Band Intensities for β-actin only. (B) Band intensities for eGFP only.
